# Weak Coupling Between Intracellular Feedback Loops Explains Dissociation of Clock Gene Dynamics

**DOI:** 10.1101/542852

**Authors:** Christoph Schmal, Daisuke Ono, Jihwan Myung, J. Patrick Pett, Sato Honma, Ken-Ichi Honma, Hanspeter Herzel, Isao T. Tokuda

## Abstract

Circadian rhythms are generated by interlocked transcriptional-translational negative feed-back loops (TTFLs), the molecular process implemented within a cell. The contributions, weighting and balancing between the multiple feedback loops remain debated. Dissociated, free-running dynamics in the expression of distinct clock genes has been described in recent experimental studies that applied various perturbations such as slice preparations, light pulses, jet-lag, and culture medium exchange. In this paper, we provide evidence that this “presumably transient” dissociation of circadian gene expression oscillations may occur at the single-cell level. Conceptual and detailed mechanistic mathematical modeling suggests that such dissociation is due to a weak interaction between multiple feedback loops present within a single cell. The dissociable loops provide insights into underlying mechanisms and general design principles of the molecular circadian clock.

## 1 Introduction

Circadian clocks are omnipresent in almost all living organisms as a consequence of adaptation to 24 h environmental fluctuations, leading to convergent evolution across different kingdoms of life [1]. Interlocked transcriptional-translational feedback loops (TTFLs) have been identified as a common design principle for the generation of intracellular rhythms. A single negative feedback is a process, in which a gene product suppresses its own expression with a time delay. Interlocking between multiple loops may have both negative and positive effects on gene expressions. In mammals, the negative feedback system that is often considered as “primary-loop” [2] consists of the *Period* (*Per1, −2, −3*) and *Chryptochrome* (*Cry-1, −2*) as well as the bHLH-PAS transcription factors *Bmal1* (also *Arntl* or *Mop3*) and *Clock*. Heterodimers of CLOCK and BMAL1 proteins enhance the transcription of *Per* and *Clock* genes by binding to their E-box promoter elements. The products of these genes, PER and CLOCK proteins, antagonize the activatory effects of the CLOCK-BMAL1 heterodimers and thus close the delayed negative feedback loop. This feedback loop will be hereinafter referred to as the *Per loop*. In addition to the “primary-loop”, a nuclear receptor loop has been identified, involving *Ror* (*Ror α*, -*β*, -*γ*) as positive regulators of *Bmal1* and *RevErb* (*RevErbα*, -*β*) as negative regulators [3,4]. Like *Per* and *Cry* genes, *RevErb* and *Ror* are transcriptionally activated by heterodimers of CLOCK and BMAL1. It has been shown by computational modeling [5] and confirmed by double-knockouts [6] that this loop plays an essential role in the rhythm generation. We will refer to this additional loop as the *Bmal-Rev loop*. It has been proposed that interlocking of such multiple loops contributes to the flexibility and robustness of the circadian system [7,8].

Complementary to experimental progress, mathematical modeling made a decisive contribution towards a better understanding of the design principles and complex dynamical behavior of the molecular circadian pacemakers across diverse organisms such as cynobacteria, fungus *Neurospora crassa*, plants, and mammals [5,9–14] as well as regulatory modules downstream of the main clock [15,16]. It has been commonly assumed that interaction of feedback loops confers robustness to molecular clock oscillations through phase- and frequency-locking of all component expressions. In the terminology of dynamical systems theory, the whole clock network constitutes a limit cycle oscillator, thereby all components form a periodic orbit of period *τ,* for which small perturbations from steady state dynamics decay with a characteristic time scale, that can be surprisingly long, even longer than 24 h. In the course of such “presumably transient” dynamics, individual components of the limit cycle oscillator may dissociate and could show different instantaneous periods, amplitude modulations and phase slips as they approach their steady state oscillations.

Recent experimental evidence shows that circadian rhythms of core clock genes dissociate at least transiently under certain conditions. *In situ* hybridization of mouse SCN revealed that circadian cycles of *mPer1* expression react more rapidly than those of *mCry1* expression to an advanced lighting schedule [17]. *Per1* and *Per2* mRNA rhythms in mouse SCN have been shown to exhibit a faster re-entrainment after a 6h jet-lag phase shift compared to those of *Bmal1, RevErbα* and *Dbp* [18]. *In freely moving single-transgenic mice expressing either a Bmal1-ELuc* or a *Per1-luc* reporter construct, re-entrainment to a new stable phase occurs at different time scales for two clock components *Bmal1* and *Per1* after application of a 9h light pulse at circadian time (CT, using the endogenous period *τ* as a reference) of 11.5h (*i.e.*, half an hour before subjective night) [19]. In addition to the behavioral studies, a dissociation of clock gene expressions has been observed among organotypic SCN slices. Measurements of bioluminescence signals in SCN slices carrying a single luciferase reporter construct revealed a significantly longer circadian period in PER2::LUC oscillations compared to *Bmal1-ELuc*, while the donor animals had identical locomotor activity periods [20]. From double-transgenic mice carrying both *Bmal1-ELuc* and *Per1-luc* reporter constructs, diverging phases were observed between the two differently colored luciferase reporters in the same slice. This leads again to significantly shorter *Bmal1-ELuc* periods across a three week long-term recording [19]. Dissociation of the two genes was observed in different reporter constructs that express luciferases with more distinct emission wavelengths [21]. Furthermore, phase response dynamics to timing of medium exchange were found to be different in *Bmal1-ELuc* and *Per2-SLR2* oscillations in cultured slices of the SCN [21]. While *Bmal1* showed a significant response to the medium exchange in neonatal mice [22], *Per2* did not [21].

Despite these experimental observations, existing mathematical models have not taken into account such long-transient dissociation of clock genes, presumably involved in different feedback loops. We use data-driven conceptual and contextual modeling approaches to identify intracellular network topologies and parameter realizations that enable experimentally observed dissociation dynamics. Our theoretical model raises new questions on the design principles of interlocked molecular loops and proposes a possibility that the biological systems may utilize such dissociation of multiple feedback loops to differential responses to external environment.

## 2 Results

### 2.1 Surrogate data analysis suggests a dissociation of *Per1* and *Bmal1* dynamics at the single cell level

*Bmal1* and *Period* (*Per1, Per2*) clock genes have been shown to exhibit differential dynamics after perturbations such as light-pulses, jet-lag, *ex vivo* slice preparations and culture medium exchange. Although their dissociation has been suggested more directly by recent experiment [19], it remains unclear whether this dynamical dissociation occurs within an intra-cellular level. To interpret the experiment of [19], two hypotheses can be considered. The first hypothesis 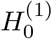 states that the dissociation takes place within a single cell. The second hypothesis 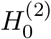, on the other hand, assumes existence of two groups of cells, in which either Bmal1 or Per1 signal is predominant. In order to examine the two hypotheses, artificial time lapse movies, *i.e.*, surrogate data [23], have been created based on either of the two hypotheses and their oscillatory properties were further analyzed. Detailed procedure for generating the surrogate data can be found in Section *Materials and Methods*. Supplementary Figure S1 illustrates various steps to generate the artificial time lapse movies.

A pixel-wise analysis of oscillatory properties in the surrogate movie data reveals qualitative differences between the two hypotheses. In the case of surrogate data generated under hypothesis 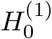, a pixel-wise comparison of Bmal1 and Per1 periods reveals two clusters in the corresponding bivariate graph, compare Figure 1 A and Supplementary Figure S2. Pixel-wise time traces in cluster 1 have a non-circadian period close to zero in both, the Bmal1 and Per1 signals. This corresponds to pixels where no SCN cells are located and, thus, the dominant peak in the Lomb Scargle periodogram is located at periods much shorter than the circadian. Time traces in cluster 2 contain a dominant circadian component in both, the Bmal1 and Per1 signals, corresponding to pixels where SCN cells have been present. This situation changes qualitatively in the case of surrogate data generated under hypothesis 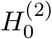, where two additional clusters emerge, see Figure 1 B. Cluster 1 still corresponds to time traces from pixels, where no significant circadian rhythm can be observed for both signals. Time traces in cluster 2 are again from pixels, where circadian periods have been detected in both Bmal1 and Per1 signals, *i.e.*, by chance a Per1 cell and a Bmal1 cell are closely located so that both signals spatially overlap with each other. In clusters 3 and 4, each pixel contains circadian component in either Bmal1 or Per1 signal, where no circadian rhythmicity is present in the other signal.

**Figure 1:**
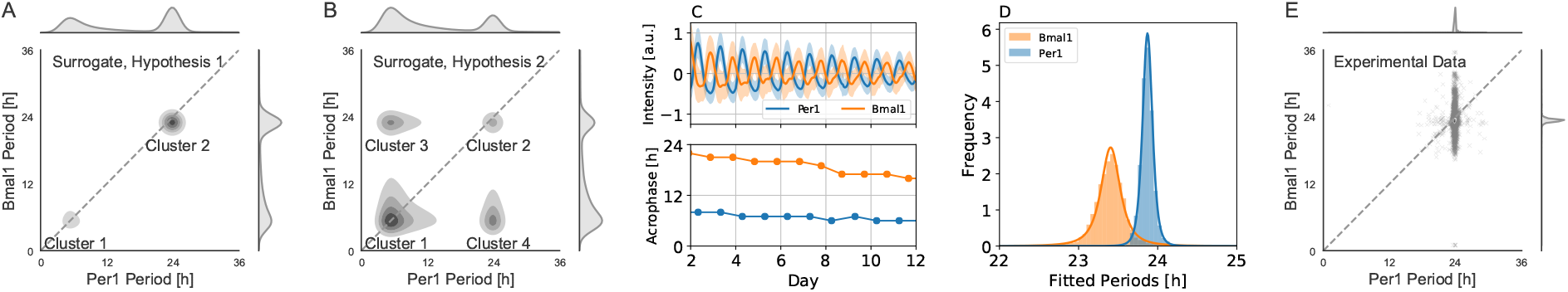
Statistical hypothesis testing indicates dissociation of *Bmal1-ELuc* and *Per1-luc* rhythms at the single cell level. A) *Gaussian* kernel density estimates in the bivariate graph of Bmal1 and Per1 oscillation periods, estimated by a Lomb Scargle analysis of surrogate time lapse movies, generated under hypothesis 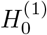, *i.e.*, dynamical dissociation at the single cell level. B) Same as panel (A) in case of hypothesis 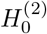, *i.e.*, randomly located cells with either Bmal1 or Per1 signal of different periods. In both panels, *N* = 150 cells have been randomly drawn. Signal intensities of 1, Bmal1 period of 23h, Per1 period of 24h, cell sizes *σ*_*G*_ = 0.0132 and noise strength of *σ*_*n*_ = 1 were used. See Supplementary Figure S1 for an example. C) *Top:* Average values (bold line) and standard deviations (shaded areas) of *Per1-luc* (blue) and *Bmal1-ELuc* (orange) signals from a cultured SCN of double transgenic mice. *Bottom:* Times of oscillation peaks (acrophases) in the averaged *Per1-luc* (blue) and *Bmal1-ELuc* (orange) signals. Compared to *Per1-luc* signals, phase drift of *Bmal1-ELuc* in oscillation peak times can be observed, suggesting a shorter *Bmal1-ELuc* period. D) Histograms of pixel-wise oscillation periods in the *Per1-luc* (blue) and *Bmal1-ELuc* (orange) signals as determined by a Lomb Scargle periodogram. Bold lines denote fits of a (non-central) Student’s t-distribution to the histogram data. The Student’s t-distribution has been preferred over normal distribution for its lower sensitivity to outliers [24]. Fitted parameters for the location (similar to the mean of a *Gaussian*) and scale parameter (similar to the standard deviation of a *Gaussian*) are ≈ 23.87 ± 0.06 h and ≈ 23.40 ± 0.13 h in case of *Per1-luc* and *Bmal1-Eluc* signals, respectively. E) Pixel-wise comparison of *Per1-luc* and *Bmal1-Eluc* periods as shown by a scatter plot (crosses) together with the corresponding kernel density estimates. The broader distribution of *Bmal1-ELuc* periods can be due to the lower SNR (signal-to-noise ratio) in comparison to the *Per1-luc* signal. Data analyzed in panels (C)-(E) correspond to the ones shown in Figure 5 of reference [19].

We next compared these qualitative features of the surrogate data with those of the corresponding experimental data. Bioluminescence recordings from *in vitro* SCN slices of neonatal double transgenic mice, expressing both *Per1-luc* and *Bmal1-ELuc* reporter constructs at the same time, have been therefore analyzed. Nearly anti-phasic relation between *Per1-luc* and *Bmal1-ELuc* oscillations can be observed in the detrended, averaged bioluminescence signals, see Figure 1 C *top*. A closer inspection of the peak times of these averaged oscillatory signals reveals a steady phase drift between *Per1-luc* and *Bmal1-ELuc* peak times, compare Figure 1 C *bottom*. This is reflected in the distributions of the pixel-wise Lomb Scargle period analysis, revealing a center of the distribution around ≈ 23.87 ± 0.06 h and ≈ 23.40 ± 0.13 h for *Per1-luc* and *Bmal1-Eluc* signals, respectively, see Figure 1 D. These distributions are in good agreement with the previously reported shorter *Bmal1-ELuc* period in comparison to *Per1-luc* [19] *or Per2-SLR2* [21] *signals in neonatal double transgenic mice. A pixel-wise comparison of Bmal1-ELuc* and *Per1-luc* oscillation periods leads to a dominant single cluster in the bivariate graph, similar to the surrogate data as generated under hypothesis 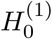, *i.e.*, dynamical dissociation at the single cell level, compare Figures 1 A and E. The broader distribution of *Bmal1-ELuc* periods can be due to its lower SNR (signal-to-noise ratio) in comparison to the *Per1-luc* signals.

To conclude our statistical hypothesis testing, comparative analysis between experimental and surrogate data supports the idea that the dissociation of *Bmal1-ELuc* and *Per1-luc* signals in slices occurs at the single cellular level (hypothesis 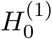).

### 2.2 A conceptual model of two weakly coupled feedback loops explains differential responses of clock gene expression upon light perturbations

In the gene-regulatory network of circadian rhythms, Bmal1 has long been thought of as a major hub. Genetic knockout of Bmal1 leads to arrhythmicity in clock gene expression and behavioral rhythms under free-running conditions [25]. However, it has been shown that constitutive expression of BMAL1 (or BMAL2) in a Bmal1^-/-^ knockout mutant mice recovers rhythmic expression of *Per2* mRNA and behavioral activity at periods similar to WT oscillations, thus questioning the necessity of rhythmic BMAL1 protein oscillations with respect to proper clock functioning [26–28]. Furthermore, computational studies suggest, that both, the negative auto-regulatory Per loop as well as the Bmal-Rev loop is able to oscillate autonomously [5,29]. Persistence of circadian rhythmicity in transgenic rats overexpressing *mPer1*, although its responsiveness to light cycles was impaired, suggests alternative feedback loop that functions without *mPer1* [30].

Motivated by these findings, we construct a conceptual model that considers interlocking of autonomously oscillating Per and Bmal-Rev intracellular feedback loops to describe transient dissociation of Bmal1 and Per1 dynamics. The dynamics of each loop is simplified by a phase oscillator, which reduces the high-dimensional limit cycle dynamics into a phase space of only a single variable, *i.e.*, the phase of oscillation *θ*. Figure 2 A illustrates the concept of phase oscillator modeling. The phase dynamics of individual loops are assumed to be governed by intrinsic angular velocities *ω*_*P*_ and *ω*_*R*_, which are related to the internal period of the Per and Bmal-Rev loop by 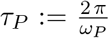 and 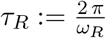, respectively, and a sinusoidal interaction function. The underlying network topology and governing equations are depicted in Figure 2 B, see also Equations (1)-(2) of Section *Material and Methods*. Parameters *K*_*R*_ and *K*_*P*_ determine the coupling strength between Per (*θ*_*P*_) and Bmal-Rev (*θ*_*R*_) loops as a function of their phase difference Δ*θ* := *θ*_*P*_ *– θ*_*R*_. The stable phase difference Δ*θ*^*^ upon complete synchronization (vanishing period difference or phase-locking) of both loops can be flexibly adjusted by parameter *β*, see Equation (6) in *Materials and Methods*. *Per1* and *Per2* transcription has been shown to exhibit acute responses to light pulses during subjective night [32–34]. We thus assume that light resetting of the core clock network solely affects the Per loop but not the Bmal-Rev loop. For the sake of simplicity, we assume a sinusoidal *Zeitgeber* signal 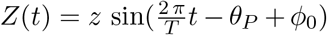, similar to previously published computational studies on entrainment of the mammalian circadian clock [35]. Here, *T* denotes the *Zeitgeber* period, while *z* determines the effective strength of the signal.

**Figure 2:**
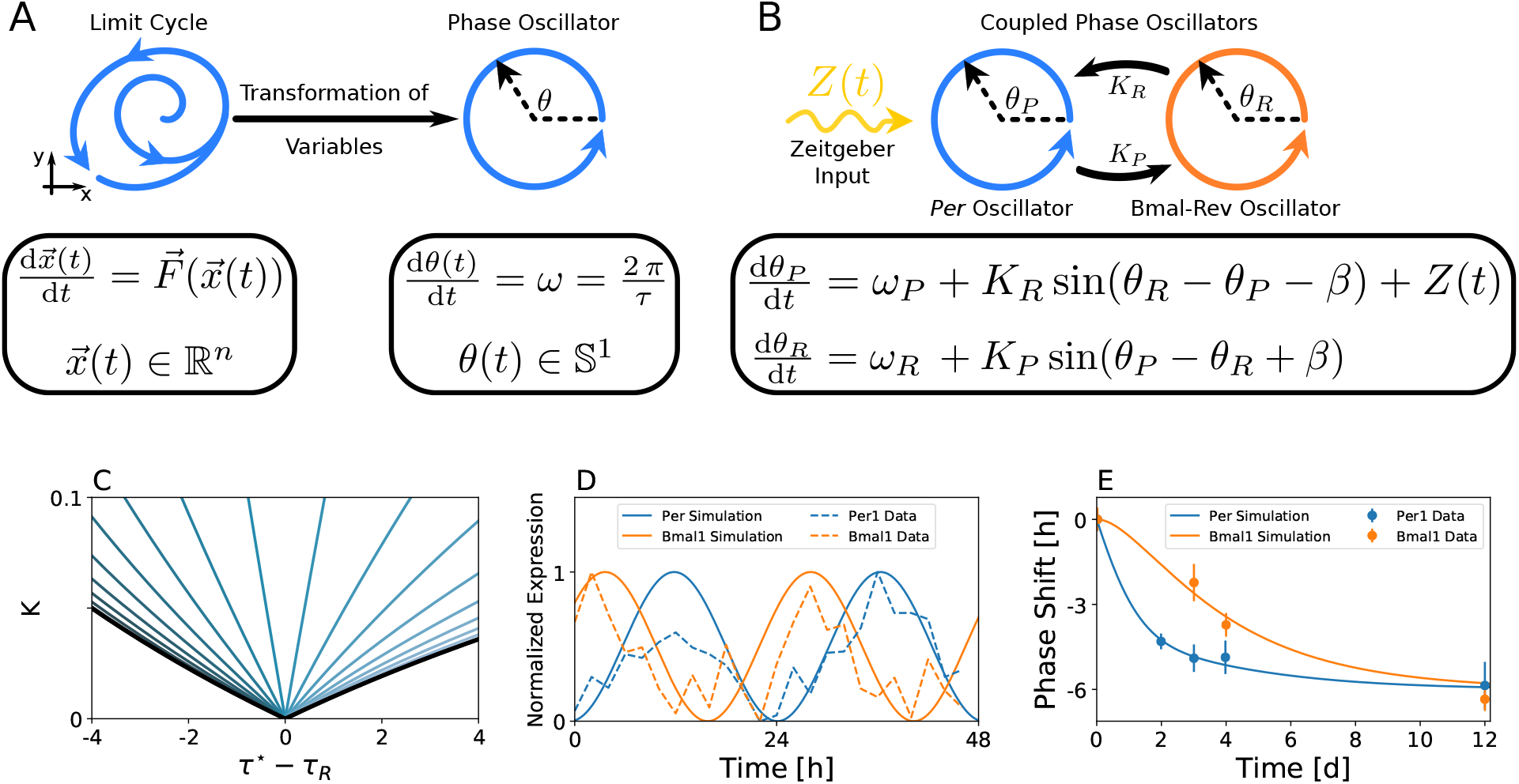
A light driven network of two coupled phase oscillators, representing the Per and Bmal-Rev loops, is able to reproduce experimental free-running and light perturbation data. A) Illustration of the phase oscillator concept. B) Schematic drawing of our conceptual model of light-driven, interlocked intra-cellular feedback loops. C) Isoclines of constant phase difference between the Per and the Bmal-Rev loops, color-coded for different values of *β*. Black lines denote the borders of synchronization between the Per and Bmal-Rev loops as determined by Equation (4) in Section *Materials and Methods*. Isoclines of constant Δ*θ*^*^ = -0.7*π*, corresponding to the experimentally observed phase difference of approximately 9 h between *Per1* and *Bmal1* expression in the time domain, are plotted and color-coded for different values of *β*, ranging from 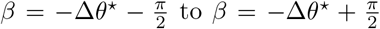 in 20 equidistant steps. The experimentally observed phase difference has been estimated by cosine fits to *Per1* and *Bmal1* circadian gene expression from high-throughput transcriptome data of 48h length at 2h sampling intervals [31], see Supplementary Figure S3. General distributions of phase differences Δ*θ*^*^ within the range of synchronization between Per and Bmal-Rev loops for different values of *β* are depicted in Supplementary Figure S4 D) Dynamics of experimentally observed *Per1* and *Bmal1* gene expression rhythms can be reproduced by the concpetual oscillator model. Bold lines denote the cosine of oscillation phases *θ*_*P*_(*t*) and *θ*_*R*_(*t*) of the corresponding Per and Bmal-Rev loop. E) Weakly coupled Per and Bmal-Rev loops can account for a faster re-entrainment of *Per1* compared to *Bmal1* gene expression oscillations after a 6h phase advancing jet-lag.

We aim to reproduce SCN expression profiles of *Per1* and *Bmal1* core clock gene oscillations under constant conditions as taken from a high-throughput transcriptome data set, recorded over a 48h period at a 2h sampling interval [31]. Under the assumption that clock genes are synchronized with a common oscillation period, we estimate a steady state phase difference of approximately 9 h or equivalently Δ*θ*^*^ ≈ *–*0.7 *π* between the *Per1* and *Bmal1* mRNA rhythms at an oscillation period of *τ* ≈ 24.53h as revealed by a cosine fit to the corresponding experimental time series, see Supplementary Figure S3. For simplicity, we assume symmetric, equally strong, coupling strength between the Per and Bmal-Rev loops (*K*_*P*_ = *K*_*R*_ =: *K*) in both coupling directions. Under constant conditions (*z* = 0), the region of phase-locking between the Per and Bmal-Rev loops (also: synchronization regime) forms a triangular shape in the parameter plane of coupling-strength (*K*) and period-detuning (*τ*\* *– τ*_*R*_). The tip of the triangle touches the point of vanishing period differences on the abscissa, see Figure 2 C and Equation (4) of Section *Materials and Methods*. Thus, for small coupling strength *K*, only small period detunings result in synchronized dynamics, while large coupling strength allows a synchronized state even for a larger detuning of periods. This is tantamount to the concept of *Arnold* tongues, describing entrainment regimes for externally forced endogenous oscillators [35,36]. For any given parameter *β* that realizes dynamics with the experimentally observed steady state phase difference Δ*θ*^*^ ≈ *–*0.7 *π* (which is given for all 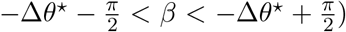, we can find an isocline of constant phase difference Δ*θ*^*^ within the synchronization regime as shown in Figure 2 C. Each pair of parameters (*K, τ*_*P*_) along such isocline gives an optimal fit to the experimentally observed phase difference as illustrated in Figure 2 D.

The sets of parameters that optimally fit *Per1* and *Bmal1* gene expression rhythms under free-running conditions can be further constrained by additionally considering the entrainment to light cycles. We therefore quantitatively compare the simulated response to a 6h advancing phase-shift in the light schedule (jet-lag) with the corresponding experimentally obtained mRNA profiles from [18], see Figure 2 E. By calculating the residual sum of squares (RSS) between simulated and experimental time series for different coupling strength *K* and *Zeitgeber* intensities *z*, we can identify for any given *β* a global optimum in the corresponding fitness landscape, see Figure 3 A. In case of *β* = 0.7 *π*, such global optimum can be found for *τ*_*P*_ ≈ 24.38h, *τ*_*R*_ ≈ 24.68h, *K* ≈ 0.043 and *z* ≈ 0.051, compare Figure 3 A. A sensitivity analysis that considers changes in the *Zeitgeber* intensity *z* and coupling strength *K* reveals that *Zeitgeber* intensity *z* mainly determines the time scale of Per response to jet-lag, while coupling strength *K* mainly determines to which extent response dynamics of the Bmal1 lags behind that of Per, compare Figures 3 B, C and Supplementary Figure S5 B, C. Generally, a larger (smaller) *Zeitgeber* intensity *z* or coupling strength *K* accelerates (decelerates) the corresponding response dynamics.

**Figure 3:**
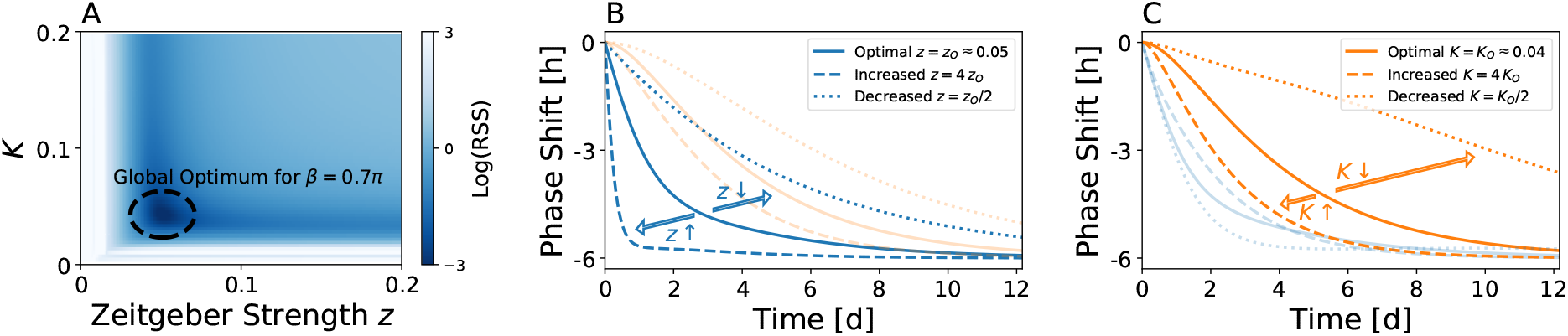
Constraining *Zeitgeber* and coupling parameters by jet-lag data. Given any *β* that allows for a reproduction of the experimentally observed phase difference between Per1 and Bmal1 oscillations under constant light conditions, the parameter (*z*) that determines the *Zeitgeber* strength can be estimated from experimental jet-lag data as demonstrated here for *β* = 0.7 *π*. A) Fitness landscape in the parameter plane of coupling constant *K* and *Zeitgeber* strength *z*. Please note that for each *K*, we assigned the parameter *ω*_*P*_ (and thus also *ω*_*R*_) along the iscoline of Figure 2 C such that the experimentally observed phase difference between Per and Bmal-Rev loops is reproduced. Colors denote the logarithm of the residual sum of squares (RSS) between the simulated and experimental jet-lag dynamics as depicted in Figure 2 E. B,C) Impact of *Zeitgeber* intensity (*z*, panel B) and coupling strength *K* between intracellular feedback loops (panel C) on jet-lag behavior of the Per (blue lines) and Bmal-Rev (orange lines) dynamics.

After application of a 9h light pulse at CT11.5h to adult mice, differential responses of *Per1-luc* and *Bmal1-ELuc* expression rhythms has been observed under *in vivo* recordings, see Figure 4 A and reference [19]. The experimentally observed phase dynamics that, after initial acute *Per1-luc* response to light, converges towards the steady phase difference Δ*θ*^*^ in the long run, can be reproduced by our conceptual model, using the “optimal” parameter set, *i.e.*, the parameter set that optimally reproduces the above described free-running and jet-lag data (for *β* = 0.7*π* as highlighted in Figure 3 A), see Figure 4 B. Again a larger (smaller) coupling strength *K* would lead to faster (slower) recovery of the steady state phase difference Δ*θ*^*^, compare the dashed (dotted) line in Figure 4 B.

**Figure 4:**
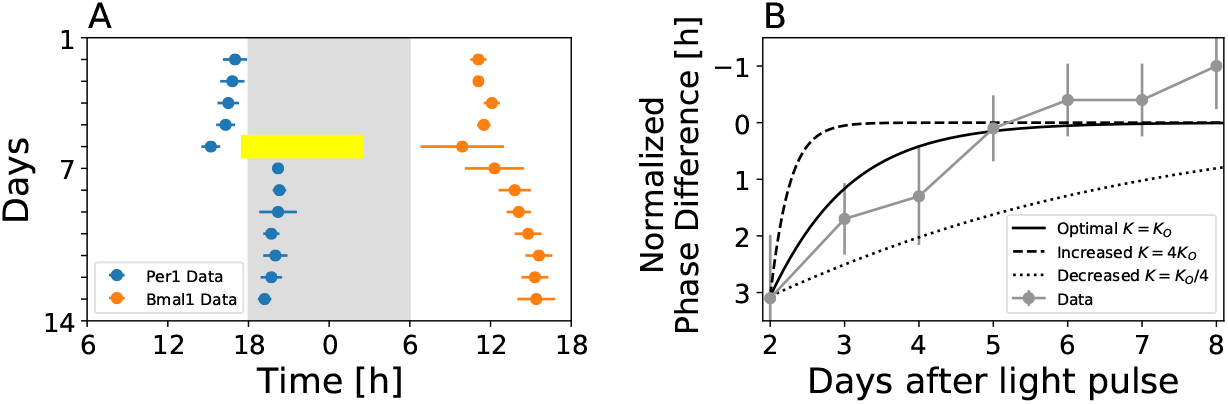
Differential response after light pulse applications depends on the coupling strength between the Per and Bmal-Rev feedback loops. A) Mean acrophases and the corresponding standard deviation of *Per1-luc* (orange) and *Bmal1-ELuc* (blue) oscillations (*n* = 3 for each reporter construct), recorded *in vivo* from single-transgenic adult mice as described in [19]. At day 5, a 9h light pulse was applied at CT11.5h (yellow bar). B) Simulated dynamics of the phase difference (Δ*θ*(*t*)) between Per and Bmal-Rev loops as given by Equation (3) of Section *Materials and Methods* for different coupling strength *K* (black lines) in comparison with the corresponding experimental data (gray). Phase differences have been normalized such that the phase difference between Per and Bmal1 oscillations one day prior to the application of the light pulse is set to zero in both, simulated and experimental time courses. Application of the light pulse leads to a perturbation from the Per1-Bmal1 free-running phase difference by approximately 3h that subsequently re-adapts within 5 days. Note that panel A is a modified reproduction of Figure 1 C in [19].

In conclusion, our conceptual model that assumes interlocking of autonomously oscillating Per and Bmal-Rev intracellular feedback loops, where only the Per loop receives direct light input, successfully describes experimental *Per1* and *Bmal1* mRNA oscillations under free-running conditions as well as dissociating dynamics upon jet-lag and 9h light pulses.

### 2.3 A minimal three-gene molecular circuit model of interlocked feedback loops successfully recapitulates free-running and light-response behavior

Can the results from our conceptual modeling be reproduced by contextual molecular circuit models that describe the network of transcriptional regulations between the core clock genes? It has been shown that condensed molecular circuit models, accounting for the interplay of cis-regulatory elements while transforming post-transcriptional regulations (*e.g.*, phosphorylation, nuclear transport, complex formation) into explicit delays, are able to faithfully reproduce experimentally observed periods and phases under free-running conditions [29]. Using a five gene model of the mammalian core oscillator network - consisting of the Bmal, Dbp, Rev, Per and Cry genes - Pett and colleagues showed that sub-networks of this model are enough to generate essential properties of the circadian oscillations, while full set of the five gene network is not needed for this purpose [37]. Such sub-modules include the auto-inhibitory regulation of *Per* and *Cry* gene expression, the Bmal-Rev loop as well as a Per-Cry-Rev repressilator motif, among others. By fitting the five gene model to clock gene expression data from 10 different tissues, it has been shown that the relative importance and balance between the sub-loops differ in a tissue-specific manner [38]. Here we aim to find a minimal molecular circuit model that accounts for the experimentally observed dynamics under free running conditions as well as dissociating dynamics of clock genes perturbed by the light input (jet-lag, light pulses).

It is known from theoretical studies that a single delayed negative feedback loop is sufficient to exhibit oscillations [29,39–41], see Figure 5 A for a schematic drawing of the corresponding network motif. By incorporating intermediate regulatory steps - such as translation, post-transcriptional or post-translational modifications - along this loop into explicit delays, we can model such single negative feedback loop by a one-variable, four parameter delay differential equation, see Equation (9) in *Materials and Methods*. For a suitable set of parameters, such one-variable model is capable of reproducing experimentally observed *Per1* gene expression rhythms under free-running conditions, see Figure 5 B. When driven by a *Zeitgeber* signal of appropriate strength (*z* = 0.21), we can mimic the response of *Per1* gene expression rhythms to a 6h phase advancing jet-lag, see Figure 5 C. Similar to the conceptual model described in the previous subsection, we incorporate the impact of light as a *Zeitgeber* signal by an additive (activatory) effect upon *Per1* transcription. Although our model was not optimized for this sake, experimentally observed entrainment phases of *Per1* mRNA [42] around midday can be reproduced by *Zeitgeber* intensity of *z* = 0.21 and model parameters as described in Section *Materials and Methods*, compare Supplementary Figure S6 A. Since the Per-one-loop model only describes the dynamics of a single clock gene, it is insufficient to reproduce the dissociating dynamics of different clock genes.

**Figure 5:**
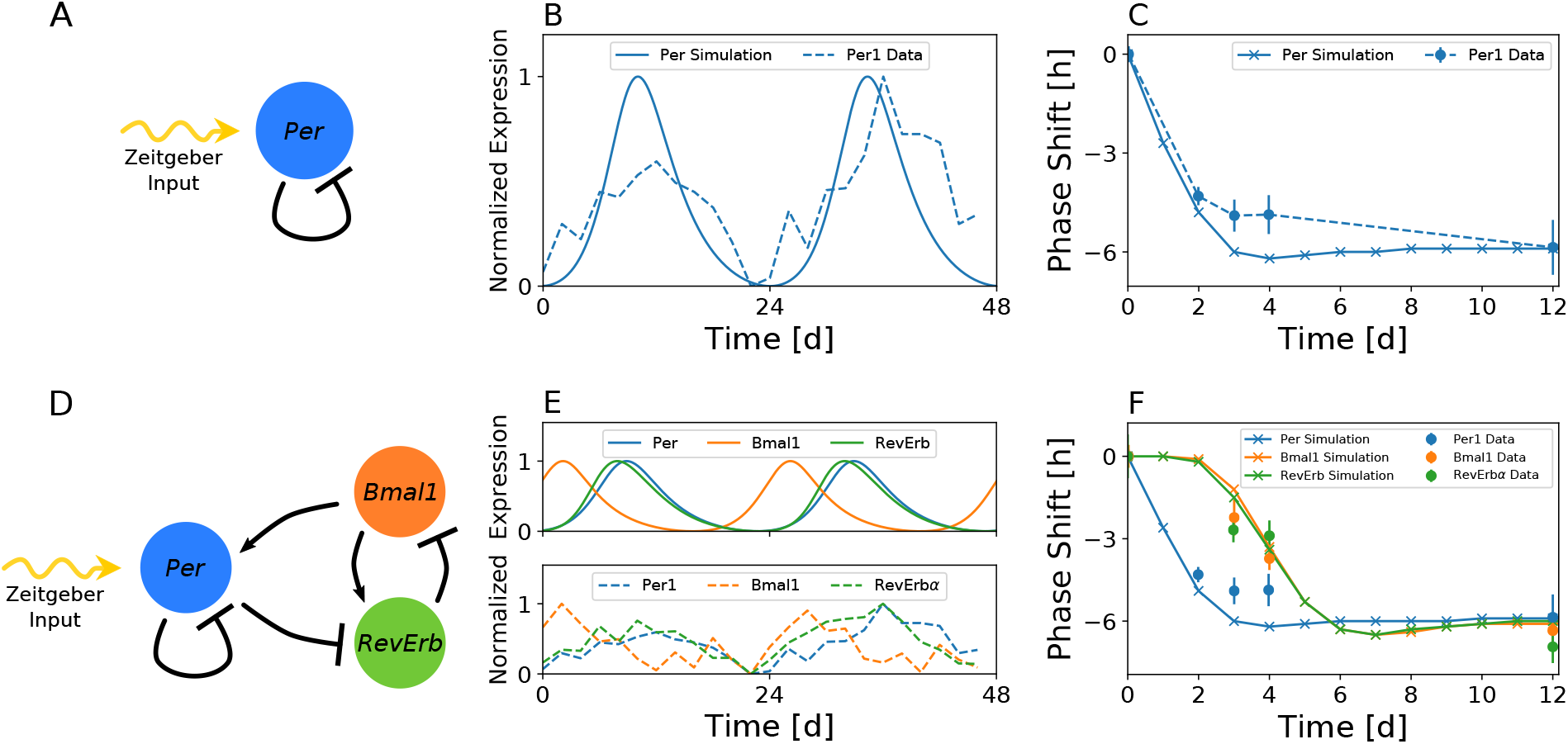
Free-running and differential jet-lag responses can be reproduced by a three-gene molecular circuit model. A) Network structure of the auto-inhibitory Per1 loop driven by light (*Zeitgeber* signal). B) The single auto-inhibitory Per1 loop is sufficient to reproduce experimentally observed Per1 gene oscillations under free running conditions for suitable sets of parameters. Simulated Per1 dynamics as well as the corresponding experimental time series from Zhang *et al.* [31] *are depicted by bold and dashed lines, respectively. C) For an appropriate Zeitgeber* intensity (*z* = 0.21), the experimentally observed Per1 mRNA response to a 6h phase advancing jet-lag can be reproduced by the light-driven single auto-inhibitory Per1 loop. D) Network structure of the light driven auto-inhibitory Per1 loop, interlocked with the Bmal-Rev loop. E) A three variable model, consisting of Per1, Bmal1 and Rev-Erb*α* genes and their transcriptional regulatory interactions is able to reproduce experimentally observed Per1, Bmal1 and Rev-Erb*α* gene expressions under free running conditions for suitable sets of parameters. F) For a suitable, intermediate strength of coupling between the Per and Bmal-Rev loops,*i.e. c*_*r*_ = 35, the experimentally observed differential response of Per1, Bmal1, and Rev-Erb*α* genes to a 6h phase advancing jet-lag can be observed. The term “intermediate” denotes strong enough coupling to synchronize the Per and Bmal-Rev loops but weak enough coupling to allow for a dissociation between them.

We therefore expand the single auto-inhibtory Per1 loop model by interlocking it with the two-gene Bmal-Rev loop, see Figure 5 D for a schematic drawing of the corresponding network motif. For a suitable set of parameters, such three-gene model is able to reproduce the experimentally observed gene expressions under free running conditions, see Figure 5 E. The extended model of interlocked Per and Bmal-Rev negative feedback loops is able to mimic the experimentally observed faster re-entrainment of *Per1* gene compared to *Bmal1* and *RevErbα* genes after a 6h phase advancing jet-lag, see Figure 5 F. As depicted in Figure 6 A and B, a relatively faster response of simulated *Per* dynamics to a 9h light pulse, applied approximately 2h after the *Per* oscillation peak under DD free-running conditions, can be observed when compared to the corresponding Bmal1 response. For a suitable set of parameters, simulated time scales of transient dynamics are in good agreement with corresponding experiment (Figure 6 A and B). As discussed in the conceptual model, response times of simulated *Bmal1* gene expression with respect to perturbations in the light schedule depend on the “coupling strength” between Per and Bmal-Rev loops. Here, the “coupling strength” can be associated with parameter *c*_*r*_ that affects the strength of inhibition of RevErb transcription by increasing the expression levels of Per gene. A smaller value of *c*_*R*_ (*i.e.*, increasing “coupling strength”) leads to a faster response of Bmal1 to a 9h light pulse, while a larger value of *c*_*R*_ (*i.e.*, decreasing “coupling strength”) slows the Bmal1 response in comparison with the nominal value of *c*_*r*_ = 35, see Figure 6 C. As the coupling between Per and Bmal-Rev feedback loops is further weakened, qualitatively different transient dynamics may emerge, in which the phase of Bmal1 moves towards a phase-advancing direction and subsequently crosses the phase of Per, see Supplementary Figure S7. The latter situation may analogously take place when Per and Bmal-Rev feedback loops are completely desynchronized, with the exception that no stable phase locking emerges after transient dynamics decayed. Thus, a complete dissociation can only be experimentally distinguished from long transient dissociating dynamics for recordings on a sufficiently long time interval.

**Figure 6:**
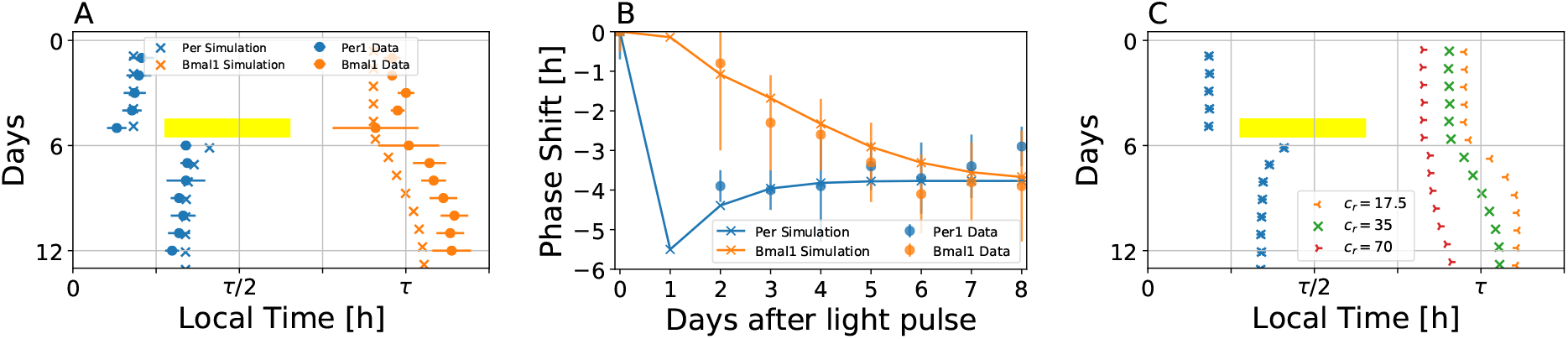
A three-gene molecular circuit model accounts for experimentally observed differential dynamics induced by a 9h light pulse. A) Simulated (crosses) and experimental (circles) acrophases of Per1 and Bmal1 gene oscillations, subject to a 9h light pulse. The yellow bar denotes the 9h light pulse in the simulated dynamics. A *Zeitgeber* intensity of *z* = 0.45 was used during the 9h light pulse. In analogy to the corresponding experimental conditions, the light pulse was applied 2.3h after the peak of Per1 expression. Note that experimental acrophase data (circles) are the same as those in Figure 4 A. The time scale represented on the x-axis has been normalized to account for different free-running periods in the light pulse experiment and the three-gene model fitted to the high throughput data. B) Dynamical evolution of the simulated (bold lines) and experimentally observed (dashed lines) phase shift of Per1 (blue) and Bmal1 (orange) genes induced by a 9h light pulse, as depicted in panel (A). C) Simulated acrophases of Per1 (blue) and Bmal1 (orange, green, red) genes, subject to a 9h light pulse, for different parameter values *c*_*r*_. Compared to the case of *c*_*R*_ = 35 depicted in panel (A), a smaller value of *c*_*R*_ = 17.5 leads to a faster response of Bmal1 to the light pulse, while a larger value of *c*_*R*_ = 70 leads to a slower response. Different values of *c*_*r*_ are highlighted by different marker symbols.

## 3 Discussions

Per1/2 and Bmal1 reporters have been routinely used in various circadian studies, where slight differences in period between the two reporters have been known. However, it has been only recently realized that these differences can be systematic and are dependent upon specific external perturbations applied to the clock system. Abrupt alterations in the *Zeitgeber* signal such as jet-lags and disruptive light pulses can lead to a dissociation of Per and Bmal1 gene expression oscillations in the SCN of live animals [17–19]. A similar dissociation has been observed after preparation of SCN slices, *i.e.*, after transferring the clock system from *in vivo* to *in vitro* conditions [19,20]. Furthermore, subsequent culture medium exchanges elicit differential phase-dependent phase shifts of *Per2* and *Bmal1* oscillations [21]. These data have been recorded at the SCN tissue level. The question remains whether the observed dynamical dissociation occurs at the single cell level or between disjoint subsets of the SCN neurons. Our surrogate data analysis, based on a comparison between experimentally obtained bioluminescence recordings and the corresponding *in silico* generated data, favors the assumption that the dynamical dissociation occurs at the single cell level. It should be noted, however, that the surrogate data of both hypotheses can show qualitatively similar features in case that the cell densities are high or the cell sizes are large that signals of neighboring cells overlap in individual pixels, see Supplementary Figure S2. Although our analysis is based on reasonable assumptions on the cell density, a definitive answer to the single cellular dissociation may require experiment on simultaneous measurements of *Per* and *Bmal1* gene expressions in isolated or sufficiently dispersed cells.

Ubiquity of the circadian rhythms in broad biological processes and organisms suggests that, despite the common underlying mechanism of negative feedback loop, there can be diverse biological implementations [43]. In mammals, more than a dozen clock genes have been described to constitute the core clock network, including *Per, Cry, Bmal, RevErb* and *Ror* genes [44]. These genes form multiple feedback loops that have different effects on regulating their own expression. Functionally redundant loops ensure robustness, while heterogeneous combinations of negative and positive feedback loops can provide higher flexibility in oscillations than a single feedback loop, such as a broader range of tunability [45]. Similarly, heterogeneous interaction of clocks may have a wider encoding capability in the tissue-level network [46], which can be reduced to two-oscillator dynamics [47]. To elucidate the transient dissociation of clock genes, we have divided the molecular feedback loops into a Per and a Bmal-Rev loop. By means of conceptual and detailed mechanistic molecular circuit models, we could show that such dissociation at the single cell level is indeed plausible within a system of multiple interlocked feedback loops.

The conceptual phase oscillator model highlights design principles for the dissociating clock gene dynamics. Oscillation phases under free-running conditions as well as time scales of transient dissociation between the Per and Bmal-Rev loops can be realized by well balanced period differences and coupling strength between the two loops. Responses of the Per loop to perturbations in the light schedule largely depend on the effective *Zeitgeber* strength. Time scales of the transient dissociation between Per and Bmal-Rev loops, which are long enough to be analogous to “internal desynchronization,” are determined mainly by their coupling strength and their individual periods. Due to the abstract nature and generality of the phase oscillator model, which is solely based on two coupled oscillators with one entity (the Per loop) being unilaterally driven by light, the results can be transferred to interpret analogous situations in other experimental settings. Based on neurotransmitter and neuropeptide release as well as afferent and efferent connections, the SCN has been functionally divided into different sub-regions, *e.g.*, core and shell [47,48]. Neurons in the SCN core release vasoactive intestinal polypeptide (VIP) and gastrin releasing peptide (GRP) and receive most of the afferent inputs via the retinohypothalamic tract that mediates light information to the SCN. This core region is surrounded by the shell region that mainly releases arginine vasopressin (AVP) and is less dense in afferent synaptic inputs mediating photic cues. Analogously to the *Per1* -*Bmal1* dissociation in response to a 6 h advance of *Zeitgeber* cycles, neurons in the SCN core entrain faster to the phase shifts compared to those in the SCN shell [49]. Here, the core that receives photic input is functionally similar to the Per loop, while the shell corresponds to the Bmal-Rev loop. Intermediate coupling between the core and the shell, that is strong enough to allow their synchronized oscillations but weak enough to exhibit “internal desynchronization” in the process of adjustment to the new phase, may explain the data.

As a more realistic molecular circuit model, a minimalistic three-gene network has been further proposed, in which negative auto-regulatory Per loop is interlocked with the Bmal-Rev feedback loop. Regulatory interactions between the three genes have been inferred from experimentally validated interactions via cis regulatory E-box, D-box and ROR elements as proposed in [29,37, 38]. An intermediate coupling strength between the two loops, that is strong enough to exhibit synchronized oscillations but weak enough to allow for transient differential dynamics of the two, can recapitulate experimentally observed dissociating dynamics induced by jet-lags and light pulses. Previously published molecular circuit models of the mammalian circadian clock mainly focused on steady state free-running, entrainment to *Zeitgeber* cycles, and mutant behaviors [5,12,29,37,38,50–52]. We here provide a mammalian intracellular clock model that additionally accounts for the transient differential behavior of clock genes in response to light perturbations. We highlight that even a minimalistic gene regulatory network, composed of not more than three genes, is able to explain a variety of complex data sets.

The minimalistic three-gene network, composed of *Bmal1, Per*, and *RevErb* genes, represents only a subset of known mutual clock gene regulations. This structure can be interpreted as a sub-module, or motif, that embeds into a more complex network of clock gene regulations, including additional elements such as *Cry, Ror* and *Dec* genes. We therefore tested whether transient dissociating dynamics of the clock genes can be identified even in a larger core clock model as described in [37,38]. Therein, the network of 20 known clock genes has been condensed into gene regulatory interactions of five groups of genes (see Supplementary Figure S8 A and [37,38]). After re-analyzing and optimizing the solutions from [37], which were obtained by fitting to SCN-specific data sets with additional “sub-network conditions” (see Supplementary Text), we found that dissociation of *Bmal1* and *Per* oscillations is possible even within this densely connected model. This implies that the present results on dissociating dynamics of clock genes are quite general and do not depend on the complexity of intracellular gene regulatory networks.

The working hypothesis of autonomously oscillating, yet coupled, intra-cellular feedback loops has a long tradition in chronobiology. In the late 1970s, long before molecular key players of mammalian circadian rhythm generation have been identified, Pittendrigh, Daan and Berde proposed two separate coupled oscillators as a means to explain splitting of behavioral activity under constant light (LL) in *Mesocricetus auratus* [53,54]. They have been termed as morning (M) and evening (E) oscillators with respect to the timing of the corresponding activity components before the splitting. Throughout the last decades, this dual oscillator concept has been applied to interpret different kinds of circadian phenomena, including bimodal activity patterns, photoperiodic entrainment properties, after-effects and internal desynchronization [55]. In mammals, different candidate genes have been proposed to constitute such dual morning-evening oscillator system, although direct evidence for the existence of intracellular M and E oscillators is still lacking. Based on differences in free-running oscillation phases and light responses, Daan *et al.* hypothesized that *Per1* and *Cry1* may constitute a M oscillator, while *Per2* and *Cry2* act as an E oscillator within a single cell [56], a concept that was later studied computationally [57]. Nuesslein-Hildesheim *et al.* proposed a dual oscillator system as composed of light-sensitive *mPer* and light-insensitive *mCry* cycles [58]. Our model provides an alternative dual-oscillator perspective based on the light-sensitive Per negative feedback loop, interlocked with the light-insensitive Bmal-Rev feedback loop. It has been shown in [19] that phase shifting behaviors of *Per1* and *Bmal1* resemble those of the activity onset and offset in behavioral rhythms, respectively. This suggests that Per and Bmal-Rev feedback loops may explain the behaviors associated with M and E oscillators.

Circadian clock serves as an internal reference of time for activity on-set and off-set locked to the phases of day and night. While the period of day-night cycles remains fixed, the day-length varies across the seasons. Phases of the activity on-set and off-set also change through the seasons. The biological clock should, therefore, be not just a robust timekeeper of 24 h cycle but also a flexible clock that adapts to such varying photoperiods. Differential responses to light between M and E oscillators can lead to a photoperiod-dependent adjustment of their phase difference, which can ultimately explain seasonal changes in behavioral activity onset, offset and activity duration (*α*) [46,55]. Our model suggests that such flexible maintenance of time is possible within a single cell. Since seasons affect all species on the planet, it would be interesting to investigate whether transient dissociations of clock genes can be analogously observed in other non-mammalian organisms such as plants, flies, or even unicellular organisms.

## 4 Materials and Methods

### 4.1 Experimental Data

#### 4.1.1 Bioluminescence recordings

The bioluminescence recordings of transgenic mice have been obtained from reference [19]. The data from SCN slice preparations, as shown and analyzed in Figure 1, has been obtained from *in vitro* brain slice preparations of neonatal transgenic mice, expressing a *Bmal1-ELuc* and *Per1-luc* reporter construct at the same time. By using a filter wheel setup with an exposure time of 29min under each condition (*i.e.*, with and without filter), both signals have been separated such that a sampling interval of Δ*t* = 1h results for the time series of *Bmal1-ELuc* and *Per1-luc* gene expression. Acrophases of *Bmal1-ELuc* and *Per1-luc* reporter constructs before and after a 9h phase light pulse, as shown in Figures 4 and 6, have been directly taken from [19]. The behavioural data have been obtained from *in vivo* optical-fiber recordings of the SCN in single transgenic freely moving adult mice, expressing either a *Bmal1-ELuc* or a *Per1-luc* reporter construct. Further details on the experimental protocols can be found in [19].

#### 4.1.2 Jet-lag experiments

In [18], changes in the rhythmicity of SCN clock gene expression after a 6h phase advance, following equinoctial LD12:12 entrainment, has been examined by *in situ* hybridization. SCN tissue has been hybridized with labeled anti-sense RNA a day before and at days 2, 3, 4, and 12 after the 6h phase advance at 6 time points per day at an equidistant sampling interval of 4h. A sine fit to the time dependent RNA profiles, as determined by densitometry, quantifies the phase shifting dynamics induced by the 6h phase advance. Such original phase shift data of Figure 2 from [18] has been extracted by the free online software WebPlotDigitizer [59] and further used to constrain our model parameters with respect to entrainment dynamics, see Figure 2 E and 5 F.

### 4.2 Surrogate Data

Statistical hypothesis testing was applied to the bioluminescence recordings of SCN slice, expressing both *Bmal1-ELuc* and *Per1-luc*, by the method of surrogate [23]. The surrogate *in silico* data that mimics experimentally obtained time-lapse recordings of double-luciferase bioluminescence reporter constructs has been created based on two null hypotheses. The first null hypothesis 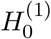 states that each of *N* single cells produces both Bmal1 and Per1 signals that differ in period, *i.e.*, Bmal1 and Per1 are assumed to be dissociated at the single cell level. The second null hypothesis 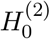 assumes that half of *N* single cells produce only a Bmal1 signal, while the other half of single cells produce only a Per1 signal with a period different from the one of the Bmal1 signal. In accordance with the experimental protocol that uses a filter wheel to seperate signals of different wavelength from *Bmal1-ELuc* and *Per1-luc* reporters, we construct a stack of two signals, the Bmal1 and Per1 signal. Spillover effects are neglected. Details of the surrogate data generation procedure are as follows: First, we locate positions of *N* single cells randomly from a two-dimensional uniform distribution. Second, periods for Bmal1 and/or Per1 signal are assigned to each of the the *N* cells based on the null hypothesis 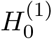 or 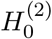. Third, Bmal1 and/or Per1 signals are generated for each cell for a total duration of 0d < *t* < 12d at a sampling rate of Δ*t* = 1h, following the experimental protocol of [19]. For the sake of simplicity, we assume that the signal *s*_*i*_(*t*), either *Bmal1-ELuc* or *Per1-luc*, in single cell *i* is described by a cosine function of maximal intensity *I*_*i*_, initial oscillation phase *ϕ*_*i*_, and period 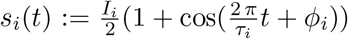. Fourth, single cell intensities of Bmal1 and Per1 signals that have been calculated at discrete positions are convoluted with a two-dimensional *Gaussian* kernel of standard deviation *σ*_*G*_ that resembles the experimentally observed size of a neuron. Subsequently, all convoluted signals are superimposed and intensity values are calculated for discrete grid positions such that the resulting grid resembles dimension of the original experimental image as well as resolution of the camera, *i.e.*, diameter of the *in silico* SCN neuron has the same dimension in units of pixels as in the corresponding experiments. Since SCN slice preparations are three-dimensional objects with neurons distributed along all three spatial dimensions, we assume *M* = 3 layers of neurons in our surrogate time lapse imaging, *i.e.*, steps 1-5 are repeated for each of the *M* layers and the signals are superimposed by assuming that the intensity drops by 50% at each layer to mimic reflection and absorption processes. Finally, observational *Gaussian* noise of zero mean and standard deviation *σ*_*n*_ is added to each grid element at each time point independently. Step-wise procedures to generate the surrogate data are illustrated in Supplementary Figure S1.

### 4.3 Time Series Analysis

Experimental data for *Per1-luc* and *Bmal1-ELuc* bioluminescence recordings of SCN slices as well as the corresponding surrogate time lapse movies are analyzed using the same custom written Python script. Time series from the experimental data are baseline detrended by means of a Hodrick Prescott filter [20], using the `hpfilter` function of the `statsmodels` Python module for a smoothing parameter 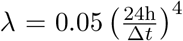 with Δ*t* being the sampling interval, as described in [60]. Oscillation periods of the detrended signals are further analyzed by a Lomb Scargle periodogram [61] in the period range of [4*h,* 48*h*] using the `lombscargle` function from the `signal` module of the `Scientific Python` package.

Additionally to the Lomb Scargle periodogram, a simple harmonic function

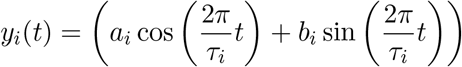

fit has been applied to the detrended time series in order to estimate the oscillatory parameters. Beside oscillation periods *τ*_*i*_, amplitudes and phases can be determined as 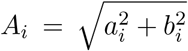 and *ϕ*_*i*_ = arctan 2(*b*_*i*_, *a*_*i*_), respectively.

### 4.4 Conceptual Model

As a conceptual model of intracellular circadian oscillation, two coupled phase oscillators [62] are constructed as follows (Figure 2 B),

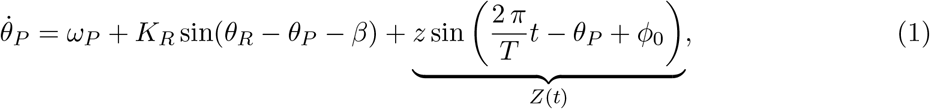

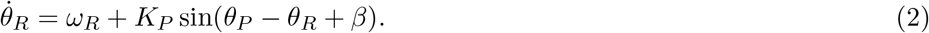

*θ*_*P*_ and *θ*_*R*_ represent oscillation phases of the Per and Bmal-Rev loops, respectively. 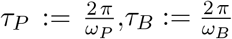 and *T* denote intrinsic periods of the Per loop, Bmal-Rev loop, and the *Zeitgeber* signal, respectively. *K*_*R*_ and *K*_*P*_ determine strength of interaction between the Per and Bmal-Rev loops, while *z* denotes strength of the light input. Parameter *β* allows for a flexible adjustment of the steady state phase difference Δ*θ* := *θ*_*P*_ *– θ*_*R*_ in the limit of vanishing frequency differences (Δ*ω* := *ω*_*P*_ *– ω*_*R*_ = 0) or infinite coupling strength (*K*_*P*_*, K*_*P*_ → ∞ for finite Δ*ω*).

Under free-running conditions (*i.e., z* = 0), Equations (1) and (2) can be rewritten as

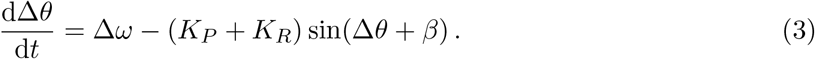

Synchronization (*i.e.*, phase-locking) between both loops, given by condition 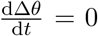, occurs for all sets of parameter that fulfill the inequality

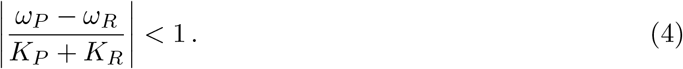

In case of such overcritical coupling or synchronization, both loops oscillate with a common angular velocity as given by the weighted arithmetic mean

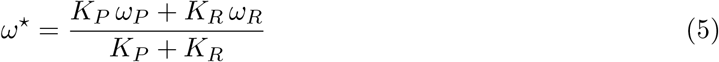

of their individual frequencies and a stable phase relationship

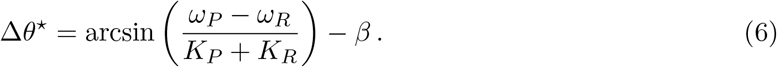

Here, the phase difference 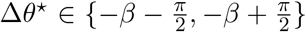 between *θ*_*P*_ and *θ*_*P*_ solely depends on *β*, the sum of the coupling strength *K*_*Σ*_ = *K*_*P*_ + *K*_*R*_ and the frequency difference Δ*ω* = *ω*_*P*_ *– ω*_*R*_.

In case of symmetric coupling (*K*_*P*_ = *K*_*R*_ =: *K*), Equation (5) is simplified to

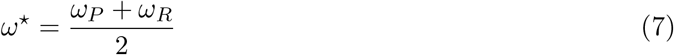

and Equation (6) can be rewritten as

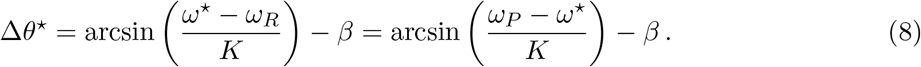

### 4.5 Detailed Mechanistic Model

Contextual molecular circuit models are developed, based on the interplay of E-box, D-Box and RRE cis-regulatory elements, as previously published [29,37,38]. While transcriptional activation and repression are described by means of (modified) Hill functions, degradation is modeled via first order kinetics. Instead of implicit delays implemented in large reaction chains, translation as well as post-transcriptional and post-translational modifications are condensed into explicit delays.

Using a previously published model [29] of Per gene expression dynamics

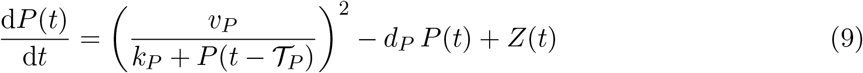

and the (modified) corresponding parameter set (*v*_*P*_ = 1, *k*_*P*_ = 0.1, *d*_*P*_ = 0.25 h^-1^, 𝒯_*P*_ = 8.333 h), we demonstrate in Figure 5 A-B that a single negative feedback loop is able to generate circadian oscillations.

In order to mimic dissociating dynamics between the Per and Bmal-Rev feedback loops, the Per single gene model of Equation (9) has been interlocked with a two-variable model, describing the Bmal-Rev negative feedback loop. The full set of Equations read as

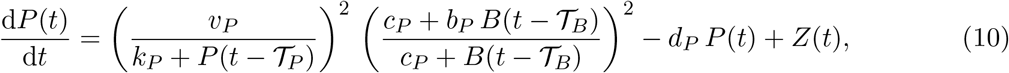

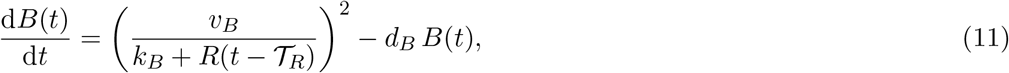

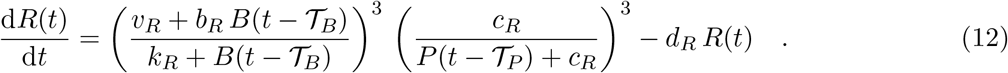

Under the assumption that light acutely induce Per transcription, time-dependent *Zeitgeber* function *Z*(*t*) appears as an additive term in Equations (9) and (10) and a square wave signal of period *T* and intensity *z* is implemented as described previously [36]. In comparison to the “full” five or six gene models of [29,37,38] that additionally include *Cry1, Ror*, and *Dbp* clock genes, here we considered only those genes and regulatory interactions that are necessary and sufficient for the occurrence of free running oscillations and entrainability of both Per and Bmal1 genes to the *Zeitgeber*. Along these lines, RevErb is a necessary network node, since the inhibitory effect of Per protein on RevErb transcription is transmitted towards Bmal1 via the inhibitory effect of RevErb on Bmal1 transcription which thus allows for light entrainment of Bmal1. Within the three node network of Per, Bmal1 and RevErb, all direct links mediated through cis regulatory elements are considered, see Figure 5 D for a schematic drawing.

Values for all parameters have been obtained from [29] and modified manually in order to adapt simulated dynamics to experimental time series data as used throughout this study. The parameter values used in our numerical simulations are *d*_*P*_ = 0.25 h^-1^, *d*_*B*_ = 0.26 h^-1^, *d*_*R*_ = 0.29 h^-1^, *v*_*P*_ = 1, *v*_*B*_ = 0.9, *v*_*R*_ = 0.6, *k*_*P*_ = 0.1, *k*_*B*_ = 0.05, *k*_*R*_ = 0.9, *c*_*P*_ = 0.1, *c*_*R*_ = 35, *b*_*P*_ = 1, *b*_*R*_ = 8, 𝒯_*P*_ = 8.333 h, 𝒯_*R*_ = 1.52 h, and 𝒯_*B*_ = 3.652 h unless otherwise stated. Hill coefficients are based upon experimentally observed binding sites as described in [29].

### 4.6 Numerics

Simulations results in Figures 2 F, 3, 4 B and Supplementary Figure S5 have been obtained by numerically solving the ordinary differential equations (1)-(3) via the `odeint` function from the `integrate` module of the `Scientific Python` package. The solutions have been drawn at equidistant intervals of Δ*t* = 0.01 h.

Simulation results from the delay differential equations (9)-(12) as seen in Figures 5, 6 and Supplementary Figures S6 and S7 have been obtained numerically by means of the `Matlab` function `dde23`, called from a Python script using the `matlab.engine` API. Again, the solutions have been drawn at equidistant intervals of Δ*t* = 0.01 h.

## Acknowledgments

This work was supported by the Japan Society for the Promotion of Science (JSPS) through grant number PE17780 and the German Research Foundation (DFG) through grant numbers HE2168/11-1 and SCHM3362/2-1. CS acknowledges support from the Joachim Herz Stiftung. JM acknowledges supports from Taiwan Ministry of Science and Technology (MOST) Grants 107-2311-B-038-001-MY2 and 107-2410-H-038-004-MY2, Taipei Medical University Grant TMU107-AE1-B15 and International Cooperation Research Plan Subsidy (TMU decree 1020004021), Taipei Medical University-Shuang Ho Hospital Collaboration Grant 107TMU-SHH-03, and Nakayama Foundation for Human Science. ITT acknowledges financial support from the JSPS (KAKENHI Nos. 16K00343, 16H05011, 17H06313, 18H02477).

## Supplementary Figures

**Figure S1:**
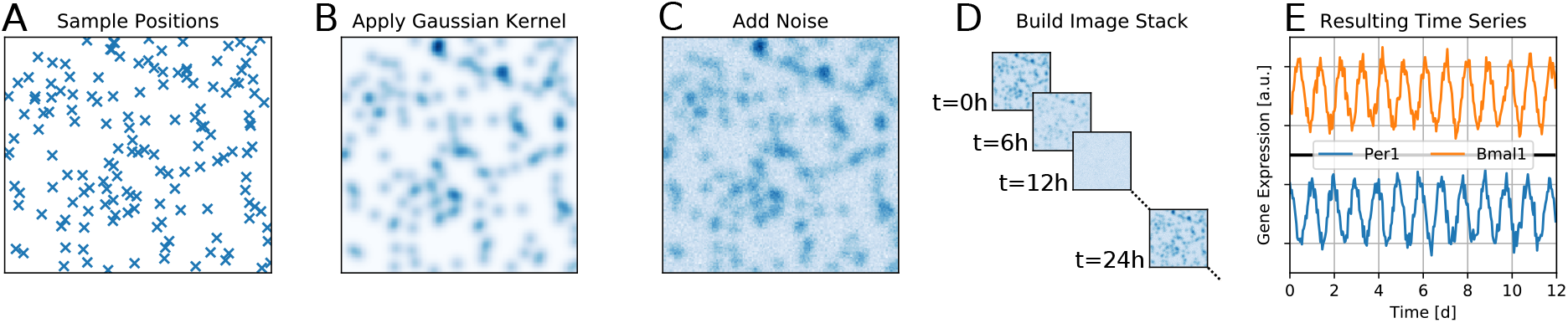
Surrogate data generation. Depicted are various steps to generate the surrogate data as described in Section *Materials and Methods* of the *Main Text*. A) *N* cells are randomly located into a square shaped space from a two-dimensional uniform distribution. B) To each cellular position, an oscillating, sinusoidal intensity signal of period *τ*_*i*_ and initial phase *ϕ*_*i*_ is assigned. To mimick the experiment, periods and initial phases of *in silico* Bmal1 or Per1 signals are set differently. At each time point *t*, the signal is convoluted with a *Gaussian* kernel of standard deviation *σ*_*G*_ in order to mimic the spatial extension of neurons. C) *Gaussian* noise of standard deviation *σ*_*n*_ is independently added to the value of each pixel, at each time point *t*. D) Illustrative sketch of the resulting surrogate data image stack for exemplary time points. E) Example of individual surrogate time series data from a single pixel for both Bmal1 (orange) and Per1 (blue) image stacks.

**Figure S2:**
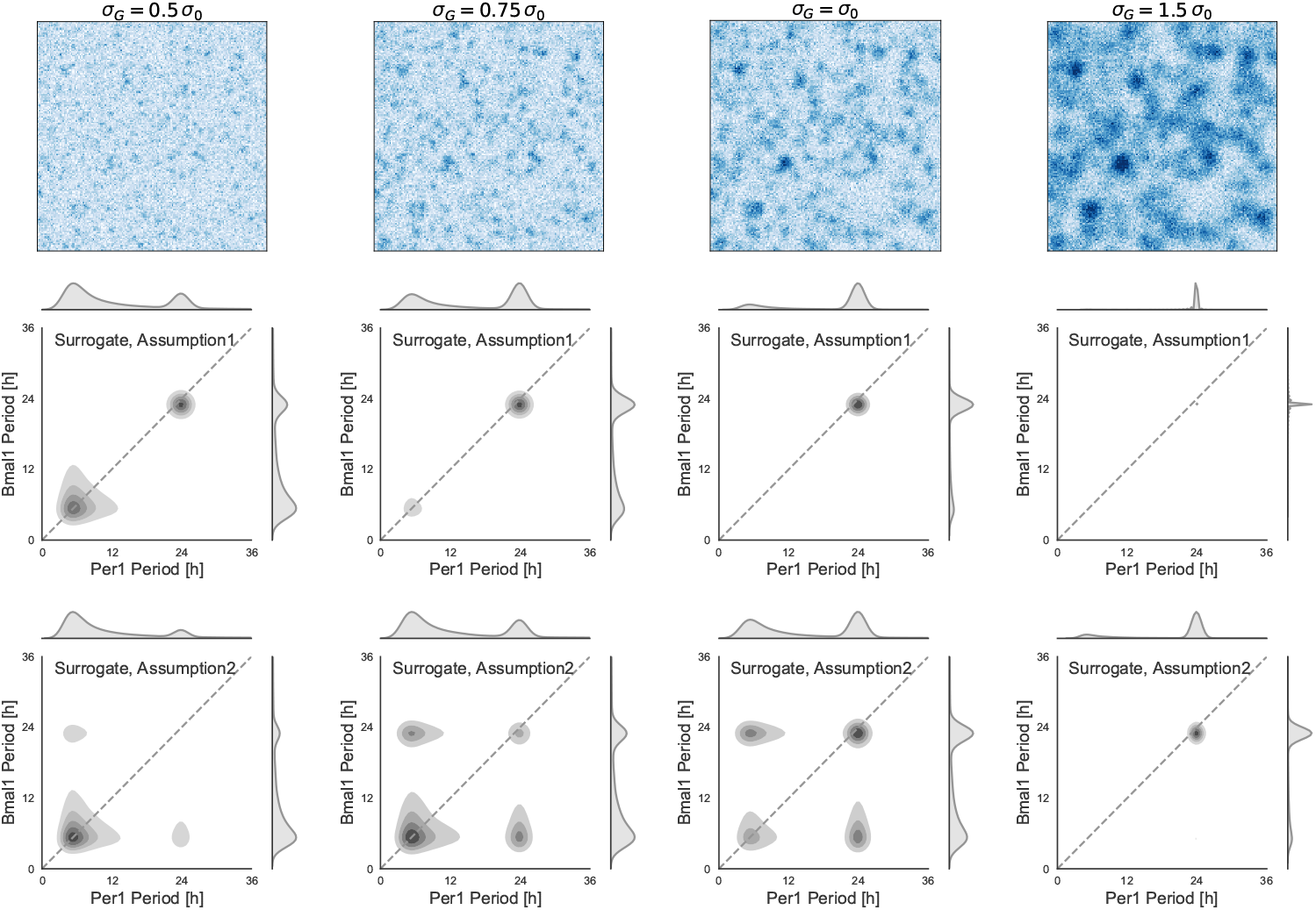
Impact of different *Gaussian* kernel width (*σ*_*G*_) on qualitative dynamical features, based on two hypotheses for surrogate data generation. *Top:* Example images of the Per1 surrogate time lapse movies at time point *t* = 0. Broadness of the *Gaussian* convolution kernels are increased from *left* to *right*, which can be associated with increasing neuron sizes or signal diffraction. Parameters *σ*_0_ = 0.0176, *N* = 150 and *σ*_*n*_ = 1 have been used. A standard deviation *σ*_*G*_ = *σ*_0_ of the *Gaussian* convolution kernel in the surrogate data generation approximates the size of an SCN neuron as recorded by the methods used in [19]. *Middle: Gaussian* kernel density estimates in the bivariate graph of Bmal1 and Per1 oscillation periods, estimated by a Lomb Scargle analysis of surrogate time lapse movies, generated under hypothesis 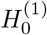, *i.e.*, dynamical dissociation at the single cell level, for an increasing *Gaussian* kernel width (*σ*_*G*_) from left to right column. *Bottom:* Same as in the *middle* panel in case of hypothesis 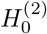, *i.e.*, randomly located cells with either a Bmal1 or Per1 signal of different periods.

**Figure S3:**
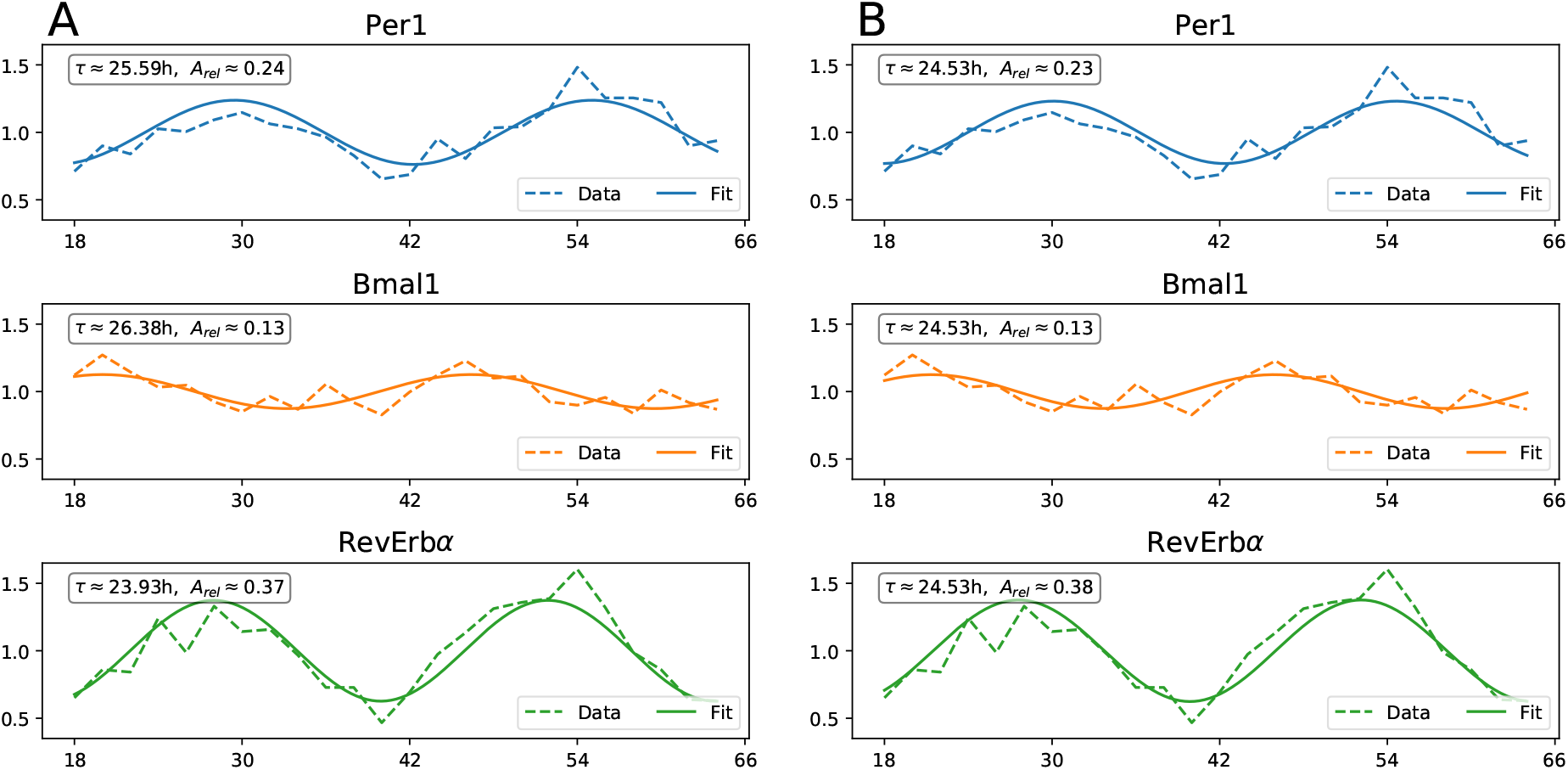
Estimation of oscillatory parameters by cosine fitting. *Bmal1, RevErbα* and *Per1* gene expression profiles of the SCN tissue data set from [31] have been first normalized by their mean expression value (such that all profiles oscillate around the value of one) and then fitted by a simple harmonic function 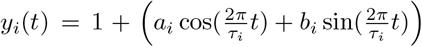. Here, indices {*i*} denote fits to different time series of the three investigated clock genes. In panel A we allow individual periods *τ*_*i*_ for all three clock genes, while in panel B we assume a synchronized state between all clock genes such that the oscillation period *τ*_*i*_ =: *τ* is shared throughout the fit to all three clock genes. The fitted relative amplitudes and phases of the individual clock gene expression rhythms are given by 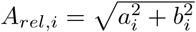 and *ϕ*_*i*_ = arctan 2(*b*_*i*_*, a*_*i*_), respectively.

**Figure S4:**
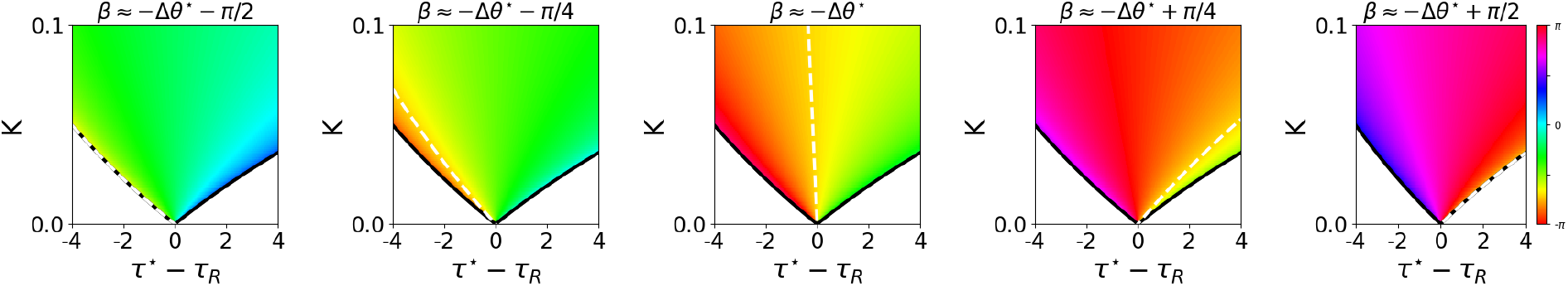
Phase differences between Per and Bmal-Rev loops in case of synchronization. Borders of synchronization (bold black lines, see Inequality (4)) and color coded phase differences (see colorbar and Equation (8)) are plotted for the conceptual phase oscillator model as given by Equations (1)-(2) of the *Main text* for different values of *β*. Δ*θ*^*^ ≈ *–*0.7 *π* denotes the experimentally observed phase differences between *Per1* and *Bmal1* gene oscillations as estimated from the SCN tissue data of [31], see also Supplementary Figure S3. Isoclines of a constant phase differences that match the experimentally observed value of Δ*θ*^*^ ≈= −0.7*π* in the *K*-(*τ*^*^ *– τ*_*R*_) parameter plane are depicted by dashed white lines. These isoclines correpsond to the color-coded isoclines of Figure 2 C of the *Main text*.

**Figure S5:**
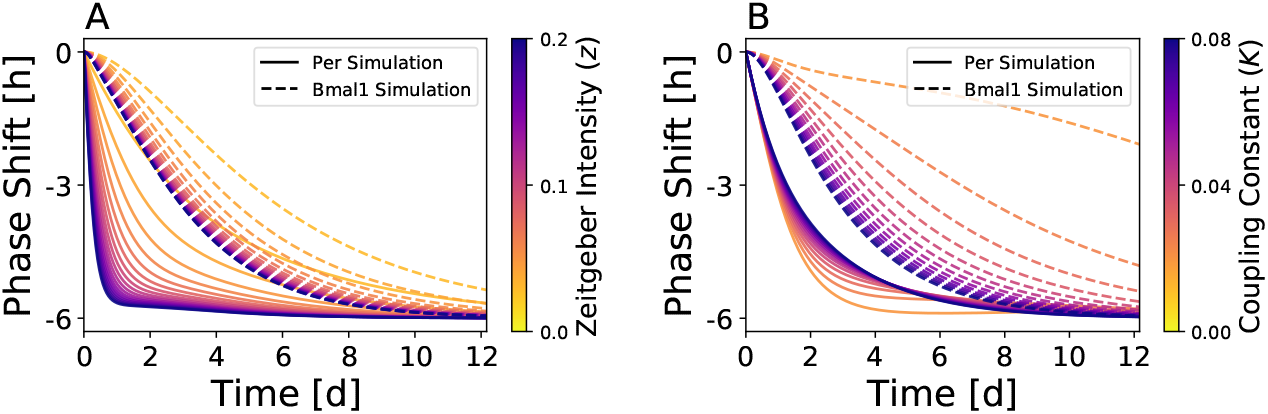
*Zeitgeber* intensity and inter-loop coupling determine jet-lag behavior. A) The Per loop dynamics shows a faster response to a 6h jet lag as the *Zeitgeber* intensity *z* is increased. Dynamics of the Bmal-Rev loop follow these dynamics although at a lower degree. B) Coupling constant *K* mainly determines how fast dynamics of the Bmal-Rev loop follow the relatively fast response of the Per loop to a 6h jet-lag. Response of the Per loop to jet-lag gets slower to some extent, since its dynamics is attracted to the Bmal-Rev loop by the symmetric coupling, which weakens the impact of *Zeitgeber* signal.

**Figure S6:**
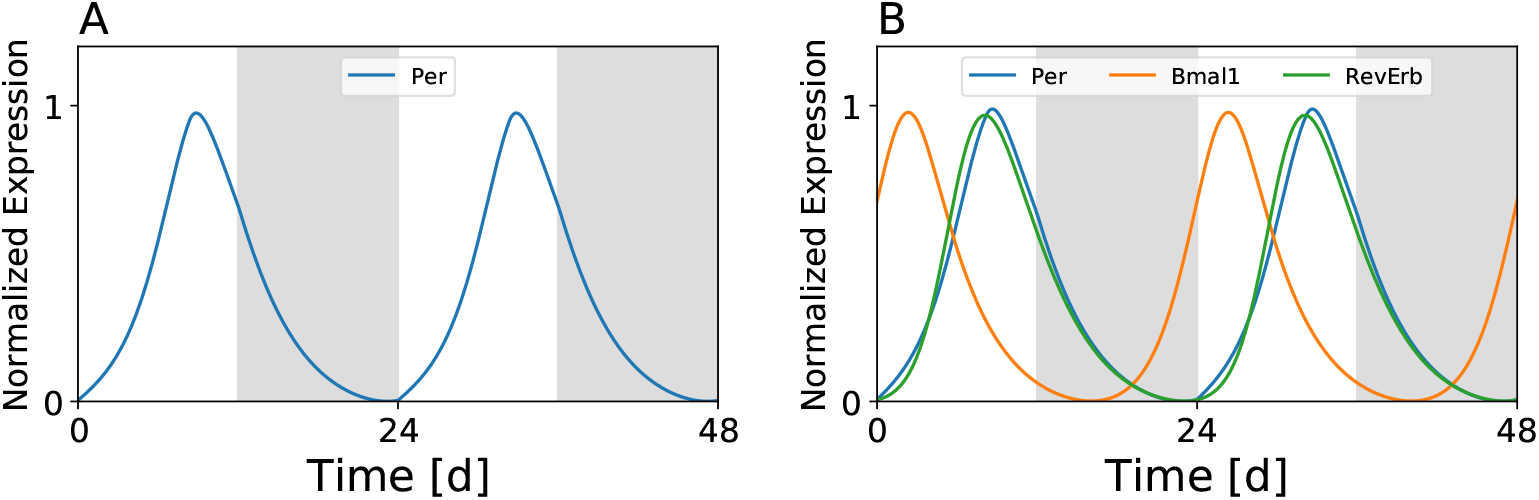
Simulated dynamics of the single- and three-gene model under entrainment conditions. A) Single-gene model. B) Three gene model. For a *Zeitgeber* intensity of *z* = 0.21 that faithfully reproduces the experimentally observed response to a 6h phase advancing jet-lag, phases of entrainment of simulated Per, Bmal1, and RevErb gene expressions qualitatively coincide with those observed in experiments. While Per and RevErb show peaks around midday, Bmal1 shows morning peaks under LD12:12 equinoctial entrainment conditions.

**Figure S7:**
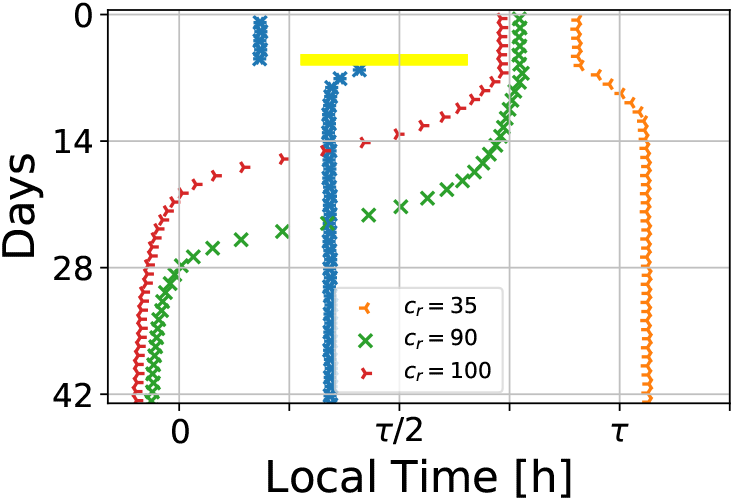
Transient dissociation for weak coupling between Per and Bmal-Rev loops in the three-gene model. Analogously to Figure 6 C of the *Main text*, simulated acrophases of Per (blue) and Bmal1 (orange, red, green) gene expressions, subject to a 9h light pulse, are depicted for different parameter values of *c*_*r*_. In case of Bmal1 oscillations, simulations with different values of *c*_*r*_ are highlighted by different marker symbols and colors. Transient dissociation dynamics can be observed for large values of *c*_*r*_ which corresponds to a weak coupling between the Per and Bmal-Rev loop due to a reduced transcriptional repression of Rev by Per.

**Figure S8:**
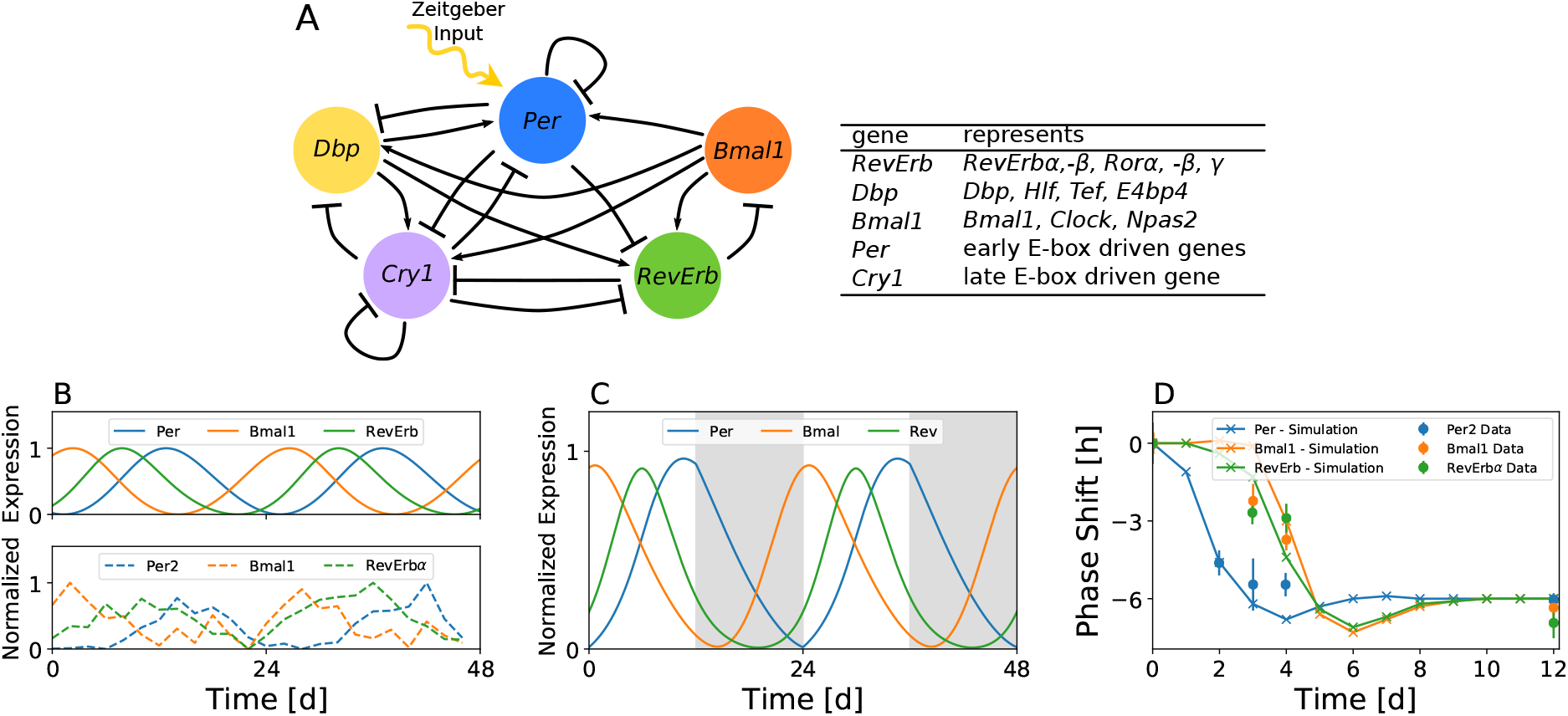
Transient dissociation can be observed within a larger mammalian core clock model. A) Schematic drawing of the regulatory core clock network. In this model, the network of 20 known clock genes has been condensed into gene regulatory interactions of five groups of genes, see table in panel (A) and references [37,38]. B) Simulation of the *Per, Bmal1*, and *RevErb* genes (top, bold lines) as well as the corresponding experimental time series (bottom, dashed lines) from SCN tissue as obtained from the high throughput study in reference [31]. Please note that kinetic parameters have been fitted to account for experimental *Per2* time series data as done in [37,38]. This results in a later phase of simulated *Per* free-running gene expressions in comparison to the conceptual phase oscillator and the three-gene model, where kinetic parameters have been optimized to account for experimental *Per1* gene expressions. C) Simulated dynamics under equinoctial LD12:12 entrainment conditions for a *Zeitgeber* intensity of *z* = 0.015. D) Simulated differential responses to a 6h phase advancing jet-lag between Per and Bmal-Rev loops together with the corresponding experimental data for *Per2, Bmal1*, and *RevErbα* genes.

## References

1. Rosbash M (2009) The implications of multiple circadian clock origins. PLoS Biol 7: e1000062.

2. Ko CH, Takahashi JS (2006) Molecular components of the mammalian circadian clock. Hum Mol Genet 15: 271–277.

3. Preitner N, Damiola F, Luis-Lopez-Molina, Zakany J, Duboule D, et al. (2002) The orphan nuclear receptor REV-ERBα controls circadian transcription within the positive limb of the mammalian circadian oscillator. Cell 110: 251–260.

4. Sato TK, Panda S, Miraglia LJ, Reyes TM, Rudic RD, et al. (2004) A functional genomics strategy reveals rora as a component of the mammalian circadian clock. Neuron 43: 527–537.

5. Relógio A, Westermark PO, Wallach T, Schellenberg K, Kramer A, et al. (2011) Tuning the mammalian circadian clock: Robust synergy of two loops. PLoS Comput Biol 7: 1–18.

6. Stratmann M, Schibler U (2012) REV-ERBs: more than the sum of the individual parts. Cell Metab 15: 791–793.

7. Akman O, Rand D, Brown P, Millar A (2010) Robustness from flexibility in the fungal circadian clock. BMC Syst Biol 4: 88.

8. Zhang EE, Kay SA (2010) Clocks not winding down: unravelling circadian networks. Nat Rev Mol Cell Bio 11: 764–776.

9. Hong CI, Jolma IW, Loros JJ, Dunlap JC, Ruoff P (2008) Simulating dark expressions and interactions of frq and wc-1 in the Neurospora circadian clock. Biophys J 94: 1221–1232.

10. Kurosawa G, Aihara K, Iwasa Y (2006) A model for the circadian rhythm of cyanobacteria that maintains oscillation without gene expression. Biophys J 91: 2015 – 2023.

11. Locke JCW, Kozma-Bognar L, Gould PD, Feher B, Kevei E, et al. (2006) Experimental validation of a predicted feedback loop in the multi-oscillator clock of Arabidopsis thaliana. Mol Syst Biol 2: 59.

12. Forger DB, Peskin CS (2003) A detailed predictive model of the mammalian circadian clock. PNAS 100: 14806–14811.

13. Thommen Q, Pfeuty B, Morant PE, Corellou F, Bouget FY, et al. (2010) Robustness of circadian clocks to daylight fluctuations: Hints from the picoeucaryote ostreococcus tauri. PLoS Comp Biol 6: e1000990.

14. De Caluwé J, Xiao Q, Hermans C, Verbruggen N, Leloup JC, et al. (2016) A Compact Model for the Complex Plant Circadian Clock. Frontiers in Plant Science 7: 74.

15. Schmal C, Reimann P, Staiger D (2013) A circadian clock-regulated toggle switch explains AtGRP7 and AtGRP8 oscillations in Arabidopsis thaliana. PLoS Comput Biol 9: e1002986.

16. Pokhilko A, Mas P, Millar AJ (2013) Modelling the widespread effects of TOC1 signalling on the plant circadian clock and its outputs. BMC Syst Biol 7: 23.

17. Reddy A, Field M, Maywood E, Hastings M (2002) Differential resynchronisation of circadian clock gene expression within the suprachiasmatic nuclei of mice subjected to experimental jet lag. Journal of Neuroscience 22: 7326–7330.

18. Kiessling S, Eichele G, Oster H (2010) Adrenal glucocorticoids have a key role in circadian resynchronization in a mouse model of jet lag. J Clin Invest 120: 2600–2609.

19. Ono D, Honma S, Nakajima Y, Kuroda S, Enoki R, et al. (2017) Dissociation of Per1 and Bmal1 circadian rhythms in the suprachiasmatic nucleus in parallel with behavioral outputs. PNAS 114: E3699–E3708.

20. Myung J, Hong S, Hatanaka F, Nakajima Y, De Schutter E, et al. (2012) Period coding of Bmal1 oscillators in the suprachiasmatic nucleus. J Neurosci 32: 8900–8918.

21. Nishide S, Honma S, Honma Ki (2018) Two coupled circadian oscillations regulate Bmal1-ELuc and Per2-SLR2 expression in the mouse suprachiasmatic nucleus. Sci Rep 8: 14765.

22. Nishide S, Honma S, Honma K (2008) The circadian pacemaker in the cultured suprachiasmatic nucleus from pup mice is highly sensitive to external perturbation. Eur J Neurosci 27: 2686–2690.

23. Theiler J, Eubank S, Longtin A, Galdrikian B, Farmer JD (1992) Testing for nonlinearity in time series: the method of surrogate data. Physica D: Nonlinear Phenomena 58: 77–94.

24. Kruschke JK (2013) Bayesian estimation supersedes the T test. J Exp Psychol Gen 142: 573–588.

25. Bunger MK, Wilsbacher LD, Moran SM, Clendenin C, Radcliffe LA, et al. (2000) Mop3 is an essential component of the master circadian pacemaker in mammals. Cell 103: 1009–1017.

26. McDearmon EL, Patel KN, Ko CH, Walisser JA, Schook AC, et al. (2006) Dissecting the functions of the mammalian clock protein BMAL1 by tissue-specific rescue in mice. Science 314: 1304–1308.

27. Liu AC, Tran HG, Zhang EE, Priest AA, Welsh DK, et al. (2008) Redundant function of REV-ERBα and β and non-essential role for Bmal1 cycling in transcriptional regulation of intracellular circadian rhythms. PLoS Genet 4: e1000023.

28. Shi S, Hida A, McGuinness OP, Wasserman DH, Yamazaki S, et al. (2010) Circadian clock gene Bmal1 is not essential; functional replacement with its paralog, Bmal2. Curr Biol 20: 316–321.

29. Korenčič A, Bordyugov G, Košir R, Rozman D, Goličnik M, et al. (2012) The interplay of cis-regulatory elements rules circadian rhythms in mouse liver. PLoS ONE 7: e46835.

30. Numano R, Yamazaki S, Umeda N, Samura T, Sujino M, et al. (2006) Constitutive expression of the period1 gene impairs behavioral and molecular circadian rhythms. PNAS 103: 3716–3721.

31. Zhang R, Lahens NF, Ballance HI, Hughes ME, Hogenesch JB (2014) A circadian gene expression atlas in mammals: implications for biology and medicine. PNAS 111: 16219–16224.

32. Shigeyoshi Y, Taguchi K, Yamamoto S, Takekida S, Yan L, et al. (1997) Light-induced resetting of a mammalian circadian clock is associated with rapid induction of the mPer1 transcript. Cell 91: 1043–1053.

33. Albrecht U, Sun ZS, Eichele G, Lee CC (1997) A differential response of two putative mammalian circadian regulators, mPer1 and mPer2, to light. Cell 91: 1055–1064.

34. Shearman LP, Zylka MJ, Weaver DR, Kolakowski LF, Reppert SM (1997) Two period homologs: Circadian expression and photic regulation in the suprachiasmatic nuclei. Neuron 19: 1261–1269.

35. Bordyugov G, Abraham U, Granada A, Rose P, Imkeller K, et al. (2015) Tuning the phase of circadian entrainment. J Royal Soc Interface 12: 20150282.

36. Schmal C, Myung J, Herzel H, Bordyugov G (2015) A theoretical study on seasonality. Front Neurol 6: 94.

37. Pett JP, Korenčič A, Wesener F, Kramer A, Herzel H (2016) Feedback loops of the mammalian circadian clock constitute repressilator. PLoS Comput Biol 12: e1005266.

38. Pett JP, Kondoff M, Bordyugov G, Kramer A, Herzel H (2018) Co-existing feedback loops generate tissue-specific circadian rhythms. Life Science Alliance 1: e201800078.

39. Goodwin BC (1965) Oscillatory behavior in enzymatic control processes. Adv Enzyme Regul 3: 425–437.

40. Scheper T, Klinkenberg D, Pennartz C, van Pelt J (1999) A mathematical model for the intracellular circadian rhythm generator. J Neurosci 19: 40–47.

41. Lema MA, Golombek DA, Echave J (2000) Delay model of the circadian pacemaker. J Theor Biol 204: 565–573.

42. Challet E, Poirel VJ, Malan A, Pévet P (2003) Light exposure during daytime modulates expression of Per1 and Per2 clock genes in the suprachiasmatic nuclei of mice. J Neurosci Res 72: 629–637.

43. Bell-Pedersen D, Cassone VM, Earnest DJ, Golden SS, Hardin PE, et al. (2005) Circadian rhythms from multiple oscillators: lessons from diverse organisms. Nat Rev Genet 6: 544–556.

44. Ukai H, Ueda HR (2010) Systems biology of mammalian circadian clocks. Annu Rev Physiol 72: 579–603.

45. Tsai TYC, Yoon SC, Ma W, Pomerening JR, Tang C, et al. (2008) Robust, tunable biological oscillations from interlinked positive and negative feedback loops. Science 321: 126–139.

46. Myung J, Hong S, DeWoskin D, De Schutter E, Forger DB, et al. (2015) GABA-mediated repulsive coupling between circadian clock neurons in the SCN encodes seasonal time. PNAS 112: 3920–3929.

47. Myung J, Pauls SD (2018) Encoding seasonal information in a twooscillator model of the multioscillator circadian clock. Euro J Neurosci 48: 2718–2727.

48. Silver R (2018) Suprachiasmatic nucleus anatomy, physiology, and neurochemistry. Oxford Research Encyclopedia of Neuroscience.

49. Nagano M, Adachi A, Nakahama Ki, Nakamura T, Tamada M, et al. (2003) An abrupt shift in the day/night cycle causes desynchrony in the mammalian circadian center. J Neurosci 23: 6141–6151.

50. Leloup JC, Goldbeter A (2003) Toward a detailed computational model for the mammalian circadian clock. PNAS 100: 7051–7056.

51. Becker-Weimann S, Wolf J, Herzel H, Kramer A (2004) Modeling feedback loops of the mammalian circadian oscillator. Biophys J 87: 3023–3034.

52. Geier F, Becker-Weimann S, Kramer A, Herzel H (2005) Entrainment in a model of the mammalian circadian oscillator. J Biol Rhythms 20: 83–93.

53. Daan S, Pittendrigh CS (1976) A functional analysis of circadian pacemakers in nocturnal rodents V. Pacemaker structure: A clock for all seasons. J Comp Physiol 106: 333–355.

54. Daan S, Berde C (1978) Two coupled oscillators: Simulations of the circadian pacemaker in mammalian activity rhythms. J Theor Biol 70: 297–313.

55. Helfrich-Förster C (2009) Does the morning and evening oscillator model fit better for flies or mice? J Biol Rhythms 24: 259–270.

56. Daan S, Albrecht U, Van Der Horst GT, Illnerová H, Roenneberg T, et al. (2001) Assembling a clock for all seasons: Are there M and E oscillators in the genes? J Biol Rhythms 16: 105–116.

57. Shiju S, Sriram K (2017) Hypothesis driven single cell dual oscillator mathematical model of circadian rhythms. PLoS ONE 12: e0177197.

58. Nuesslein-Hildesheim B, O’brien JA, Ebling FJP, Maywood ES, Hastings MH (2000) The circadian cycle of mPER clock gene products in the suprachiasmatic nucleus of the Siberian hamster encodes both daily and seasonal time. Eur J Neurosci 12: 2856–2864.

59. Rohatgi A (2000). WebPlotDigitizer, available on https://automeris.io/webplotdigitizer.

60. Schmal C, Herzog ED, Herzel H (2018) Measuring relative coupling strength in circadian systems. J Biol Rhythms 33: 84–98.

61. Ruf T (1999) The Lomb-Scargle periodogram in biological rhythm research: Analysis of incomplete and unequally spaced time-series. Biol Rhythm Res 30: 178–201.

62. Kuramoto Y (1984) Chemical oscillations, waves, and turbulence. Springer, Berlin.

