## Supplementary Text for "Weak Coupling Between Intracellular Feedback Loops Explains Dissociation of Clock Gene Dynamics"

### *Dissociation of oscillators in a large core clock model*

#### Previous model versions

By repeatedly fitting a core clock model [1] to tissue-specific gene expression data sets [2], *Pett et al.* obtained a variety of model versions with physiologically reasonable parameter settings [3]. The composition of oscillators constituted by sub-networks of these models was analyzed in detail, often showing synergies of several co-existing loops in each fit.

The model versions obtained for SCN often include *Bmal1* and *Per* oscillators. However, they generally do not appear to generate rhythms independently, such that stopping one oscillator disrupts rhythms completely. This is not unexpected though, since the criterion for optimization was solely the reproduction of experimental time series features, but not the existence of independent oscillators. Nevertheless, small parameter changes could already render the *Bmal1* and *Per* oscillators independent, such that they can mutually compensate for a disruption—or possibly generate dissociating rhythms.

#### Refitting core clock models with additional conditions

We adjusted the SCN-specific model versions by including criteria for independent oscillators. Models were re-optimized using existing SCN-specific parameter sets from [3] as starting conditions. The same methods were used as described in [3], with the exception that *Bmal1* and *Per* sub-networks were specifically required to generate rhythms. To this end, the clamping-strategy of *Pett et al.* was employed in an additional part of the scoring function. For both oscillators necessary and sufficient conditions of rhythm generating capability were tested, resulting in total in four additionally tested sub-models.

For example, in case of the *Bmal-Rev* oscillator, a clamped version was simulated in which only the regulations corresponding to its feedback mechanism were active (i). The sufficient condition of rhythm generation for this oscillator was ensured by requiring oscillation of this sub-model in the optimization. In addition, the generation of rhythms by this oscillator was ensured in a setting, in which the necessary conditions of all other oscillators where precluded in a minimal way (ii). This was achieved by clamping a single regulation of each feedback mechanism corresponding to the oscillators. Requiring rhythms of this sub-model ensured that the *Bmal-Rev* oscillator generated rhythms in a setting minimally altered from the default situation. Optimization including both types of sub-models was necessary to obtain reliable results with independent oscillators in the final model versions. An illustration of the four tested sub-models is shown in Figure **ST-1**.

While the same oscillation features of the time courses as in [3] were optimized for the full model, only the period and fold change of oscillations were optimized for the sub-models. Since sub-models represent anyway a disrupted version of the real core clock, however, their scores were only weighted 25% compared to the score of the complete model. Employing this modification of the scoring function, we were able to control which mechanisms generate oscillations in the designed models, while still resembling the data.

In order to favor oscillator dissociation in our adjusted model versions, we included a further modification of the scoring function: Strong regulations connecting the two oscillators were punished. Weaker regulations correspond to a smaller coupling strength and dissociation may therefore occur more easily.

The scoring function for the full model is given by Equation **ST-1**:

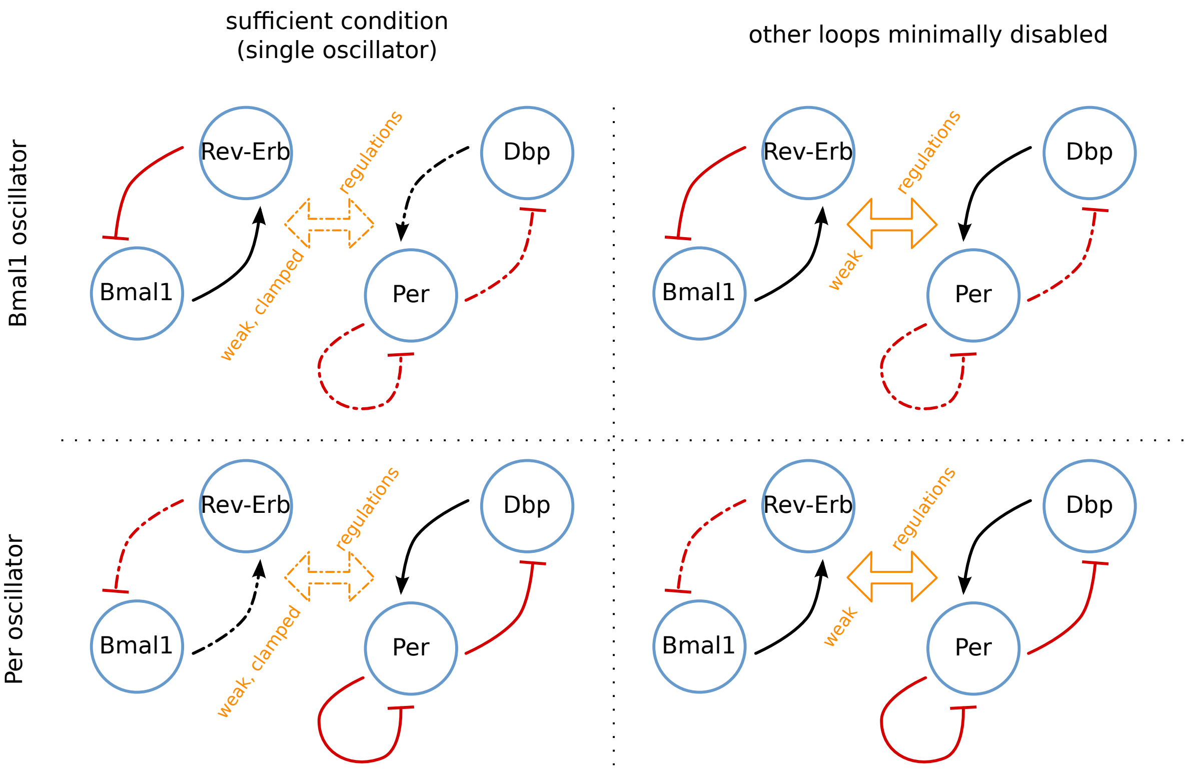


**Figure ST-1**: Tested sub-models: solid and interrupted lines correspond to active and clamped regulations, respectively. For each oscillator a model was scored in which only the corresponding loops were active. In addition, for the same oscillator another model was scored with other negative loops disrupted in a minimal way.

**Equation ST-1** Scoring function of full model

$$score_{full} = \frac{(period_{sim} - period_{exp})^2}{{tol_{period}}^2} + \sum \frac{(phase_{sim} - phase_{exp})^2}{{tol_{phase}}^2} + \sum \frac{(foldch_{sim} - foldch_{exp})^2}{{tol_{foldch}}^2} + \sum \frac{(5/param_{coupling})^2}{{tol_{coupling}}^2}$$

Where the tolerances $tol_{<.>}$ are defined as in [3] with the exception of the additional tolerance $tol_{coupling}$ for the coupling parameters which is set to 1. $param_{coupling}$ are parameters of the model that correspond to the strength of regulations connecting the two oscillators.

Furthermore, the scoring functions of the sub-models are given by Equation **ST-2**:

**Equation ST-2** Scoring functions of sub-models

$$score_{sub} = \frac{(period_{sim} - period_{exp})^2}{{tol_{period}}^2} + \sum \frac{(foldch_{sim} - foldch_{exp})^2}{{tol_{foldch}}^2}$$

The total score is then given by $score_{total} = score_{full} + 0.25 \cdot \sum score_{sub}$, where scores of four sub-models are computed as described above.

#### Equations of the large core clock model

Equations for the large core clock model are the same as in references [3] and read as

$$\frac{d\,[Bmal1]}{dt} = \underbrace{\left(\frac{1}{\frac{[\mathrm{RevErb}_\alpha](\mathrm{t}-\mathrm{del2})}{\mathrm{ar1}}+1}\right)^2}_{\text{Rev-erb-}\alpha\text{ (-)}} - \mathrm{d1}\,[Bmal1]\,(t)$$

$$\frac{d\,[RevErb_\alpha]}{dt} = \underbrace{\left(\frac{\frac{\mathrm{b\_RevErb}\,[\mathrm{Bmal1}](\mathrm{t}-\mathrm{del1})}{\mathrm{ba2}}+1}{\frac{[\mathrm{Bmal1}](\mathrm{t}-\mathrm{del1})}{\mathrm{ba2}}+1}\right)^3}_{\text{Bmal1 (+)}} \underbrace{\left(\frac{1}{\frac{[\mathrm{Per2}](\mathrm{t}-\mathrm{del3})}{\mathrm{cr2}}+1}\right)^3}_{\text{Per2 (-)}}$$
$$\underbrace{\left(\frac{\frac{\mathrm{f\_RevErb}\,[\mathrm{Dbp}](\mathrm{t}-\mathrm{del5})}{\mathrm{fa2}}+1}{\frac{[\mathrm{Dbp}](\mathrm{t}-\mathrm{del5})}{\mathrm{fa2}}+1}\right)}_{\text{Dbp (+)}} \underbrace{\left(\frac{1}{\frac{[\mathrm{Cry1}](\mathrm{t}-\mathrm{del4})}{\mathrm{gr2}}+1}\right)^3}_{\text{Cry1 (-)}} - \mathrm{d2}\,[RevErb_\alpha]\,(t)$$

$$\frac{d\,[Per2]}{dt} = \underbrace{\left(\frac{\frac{\mathrm{b\_Per2}\,[\mathrm{Bmal1}](\mathrm{t}-\mathrm{del1})}{\mathrm{ba3}}+1}{\frac{[\mathrm{Bmal1}](\mathrm{t}-\mathrm{del1})}{\mathrm{ba3}}+1}\right)^2}_{\text{Bmal1 (+)}} \underbrace{\left(\frac{1}{\frac{[\mathrm{Per2}](\mathrm{t}-\mathrm{del3})}{\mathrm{cr3}}+1}\right)^2}_{\text{Per2 (-)}} \underbrace{\left(\frac{\frac{\mathrm{f\_Per2}\,[\mathrm{Dbp}](\mathrm{t}-\mathrm{del5})}{\mathrm{fa3}}+1}{\frac{[\mathrm{Dbp}](\mathrm{t}-\mathrm{del5})}{\mathrm{fa3}}+1}\right)}_{\text{Dbp (+)}} \underbrace{\left(\frac{1}{\frac{[\mathrm{Cry1}](\mathrm{t}-\mathrm{del4})}{\mathrm{gr3}}+1}\right)^2}_{\text{Cry1 (-)}} - \mathrm{d3}\,[Per2]\,(t)$$

$$\frac{d\,[Cry1]}{dt} = \underbrace{\left(\frac{1}{\frac{[\mathrm{RevErb}_\alpha](\mathrm{t}-\mathrm{del2})}{\mathrm{ar4}}+1}\right)^2}_{\text{Rev-erb-}\alpha\text{ (-)}} \underbrace{\left(\frac{\frac{\mathrm{b\_Cry1}\,[\mathrm{Bmal1}](\mathrm{t}-\mathrm{del1})}{\mathrm{ba4}}+1}{\frac{[\mathrm{Bmal1}](\mathrm{t}-\mathrm{del1})}{\mathrm{ba4}}+1}\right)^2}_{\text{Bmal1 (+)}}$$
$$\underbrace{\left(\frac{1}{\frac{[\mathrm{Per2}](\mathrm{t}-\mathrm{del3})}{\mathrm{cr4}}+1}\right)^2}_{\text{Per2 (-)}} \underbrace{\left(\frac{\frac{\mathrm{f\_Cry1}\,[\mathrm{Dbp}](\mathrm{t}-\mathrm{del5})}{\mathrm{fa4}}+1}{\frac{[\mathrm{Dbp}](\mathrm{t}-\mathrm{del5})}{\mathrm{fa4}}+1}\right)}_{\text{Dbp (+)}} \underbrace{\left(\frac{1}{\frac{[\mathrm{Cry1}](\mathrm{t}-\mathrm{del4})}{\mathrm{gr4}}+1}\right)^2}_{\text{Cry1 (-)}} - \mathrm{d4}\,[Cry1]\,(t)$$

$$\frac{d\,[Dbp]}{dt} = \underbrace{\left(\frac{\frac{\mathrm{b\_Dbp}\,[\mathrm{Bmal1}](\mathrm{t}-\mathrm{del1})}{\mathrm{ba5}}+1}{\frac{[\mathrm{Bmal1}](\mathrm{t}-\mathrm{del1})}{\mathrm{ba5}}+1}\right)^3}_{\text{Bmal1 (+)}} \underbrace{\left(\frac{1}{\frac{[\mathrm{Per2}](\mathrm{t}-\mathrm{del3})}{\mathrm{cr5}}+1}\right)^3}_{\text{Per2 (-)}} \underbrace{\left(\frac{1}{\frac{[\mathrm{Cry1}](\mathrm{t}-\mathrm{del4})}{\mathrm{gr5}}+1}\right)^3}_{\text{Cry1 (-)}} - \mathrm{d5}\,[Dbp]\,(t) \quad .$$

#### Kinetic parameters

After re-analyzing and optimizing solutions from [3], we found several parameter sets that show a transient dissociation between simulated *Per* and *Bmal1* gene expression oscillations after a 9h light pulse or a 6h phase advancing jet-lag. Kinetic parameters found by the optimization procedure as described above have been further fine-tuned manually in order to faithfully mimic the pertinent time scales of transient dissociation dynamics. A representative parameter set that has been used to simulate dynamics as shown in Supplementary Figure S8 reads as

| del1 | del2 | del3 | del4 | del5 | d1 | d2 | d3 | d4 | d5 |
|---|---|---|---|---|---|---|---|---|---|
| 4.15 | 2.25 | 5.69 | 5.73 | 1.06 | 0.10 | 0.98 | 0.10 | 0.15 | 0.30 |

| ar1 | ar4 | cr2 | cr3 | cr4 | cr5 | gr2 | gr3 | gr4 | gr5 | b_RevErb | ba2 |
|---|---|---|---|---|---|---|---|---|---|---|---|
| 16.80 | 97.64 | 1.07 | 0.01 | 0.01 | 0.09 | 11.41 | 25.01 | 12.56 | 10.17 | 37.15 | 10.67 |

| b_Per2 | ba3 | b_Cry1 | ba4 | b_Dbp | ba5 | fa2 | f_RevErb | fa3 | d_Per2 | fa4 | d_Cry1 |
|---|---|---|---|---|---|---|---|---|---|---|---|
| 1.11 | 18.40 | 1.05 | 8.13 | 1.06 | 27.32 | 25.66 | 1.02 | 0.48 | 208.96 | 0.30 | 10.20 |

#### References


[1] Korenčič A, Košir R, Bordyugov G, Lehmann R, Rozman D, Herzel H. Timing of circadian genes in mammalian tissues. Sci Rep. 2014;4:5782.

[2] Zhang R, Lahens NF, Ballance HI, Hughes ME, Hogenesch JB. A circadian gene expression atlas in mammals: implications for biology and medicine. Proc Natl Acad Sci USA. 2014;111:16219–16224.

[3] Pett JP, Kondoff M, Bordyugov G, Kramer A, Herzel H. Co-existing feedback loops generate tissue-specific circadian rhythms. Life Science Alliance. 2018;1:e201800078.
